# Evolution of ST-4821 clonal complex hyperinvasive and quinolone-resistant meningococci: the next meningococcal pandemic?

**DOI:** 10.1101/2020.09.24.312546

**Authors:** Mingliang Chen, Odile B. Harrison, Holly B. Bratcher, Zhiyan Bo, Keith A. Jolley, Charlene M.C. Rodrigues, James E. Bray, Qinglan Guo, Xi Zhang, Min Chen, Martin C.J. Maiden

## Abstract

The expansion of quinolone-resistant *Neisseria meningitidis* clone China^CC4821-R1-C/B^ from ST-4821 clonal complex (cc4821) caused a serogroup shift from serogroup A to C in invasive meningococcal disease (IMD) in China. To establish the relationship among globally distributed cc4821 meningococci, we analysed whole genome sequence data from 173 cc4821 meningococci isolated in four continents from 1972-2019. These meningococci clustered into four sub-lineages (1-4), with sub-lineage 1 primarily comprising serogroup C IMD isolates (82%, 41/50). Most isolates from outside China formed a distinct sub-lineage (81.6%, 40/49, the Europe-USA cluster), with the typical strain designation B:P1.17-6,23:F3-36:ST-3200(cc4821) and harbouring mutations in penicillin-binding protein 2. These data show that the quinolone-resistant clone China^CC4821-R1-C/B^ has expanded to other countries. The increasing global distribution of B:cc4821 meningococci raises concern that cc4821 has the potential to cause a global pandemic and, this would be challenging to control though there is indirect evidence that Trumenba^^®^^ vaccine might afford some protection.

---

*Neisseria meningitidis*, a leading cause of bacterial meningitis and septicaemia globally, is responsible for approximately 1.2 million invasive meningococcal disease (IMD) cases annually, with a case fatality rate of 11% (*1*). Meningococci are classified into 12 serogroups based on capsular polysaccharide (*1*), while genetic relationships among isolates are defined by clonal complexes (ccs) identified by multi-locus sequence typing (MLST) which are surrogates for lineages (*2*). The relationship among serogroups, ccs (lineages), and IMD fluctuates over time and by geographic location, but IMD isolates are dominated by particular ccs (hyperinvasive lineages) usually associated with one of the six disease-causing serogroups (A, B, C, W, X, and Y).

In China, the national dissemination of hyperinvasive ST-4821 clonal complex (cc4821) meningococci led to a shift in IMD epidemiology from mostly serogroup A (MenA) to predominantly MenC (*3–4*). Although no quinolone resistance was identified in cc4821 in China in the period 1965 to 1985 (*5*), from 2005 onwards high-frequency resistance (79%) occurred, due to the expansion of the quinolone-resistant clone China^CC4821-R1-C/B^ (*5*). Previous studies discovered cc4821 can be divided into two groups, with Group 1 associated with IMD (*6–7*). Studies of a member of Group 1, isolate 053442, identified six strain-specific genome regions resulting from horizontal gene transfer (HGT) (*8*). This was consistent with the emergence of China^CC4821-R1-C/B^ and indicated that the acquisition of quinolone resistance was associated with multiple HGT events within genes encoding surface antigens (*6*), although the donors of these events were not identified.

Globally, the number of cc4821 IMD isolates has increased. When cc4821 was first identified (*4*), isolates were confined to China (Mainland China and Taiwan) (*9*); however, by the time of writing (June 2020), 59 cc4821 isolates had been identified in 19 counties worldwide (Figure 1). Moreover, three IMD cases, caused by quinolone-resistant cc4821 isolates, were reported in Canada (n=2, in 2013 and 2014) and Japan (n=1, in 2017) (*10–11*), and three other cc4821 isolates were found to colonise the anorectal tract of men who have sex with men (MSM) (*12*). Here, we investigate the genomic events leading to the emergence and expansion of hyperinvasive cc4821 meningococci by describing the phylogenetic relationships among meningococci with different serogroups (C, B, W, and nongroupable), sources (IMD, carriage, and MSM), locations (other countries vs China), and dates (1972-1978 vs 2005-2019) of isolation. We assess genes encoding key antigens genes and antimicrobial resistance phenotypes, identify putative donors of the HGT events unique to the epidemic and quinolone-resistant clone China^CC4821-R1-C/B^, and characterise isolates from other countries.

**Figure 1.**
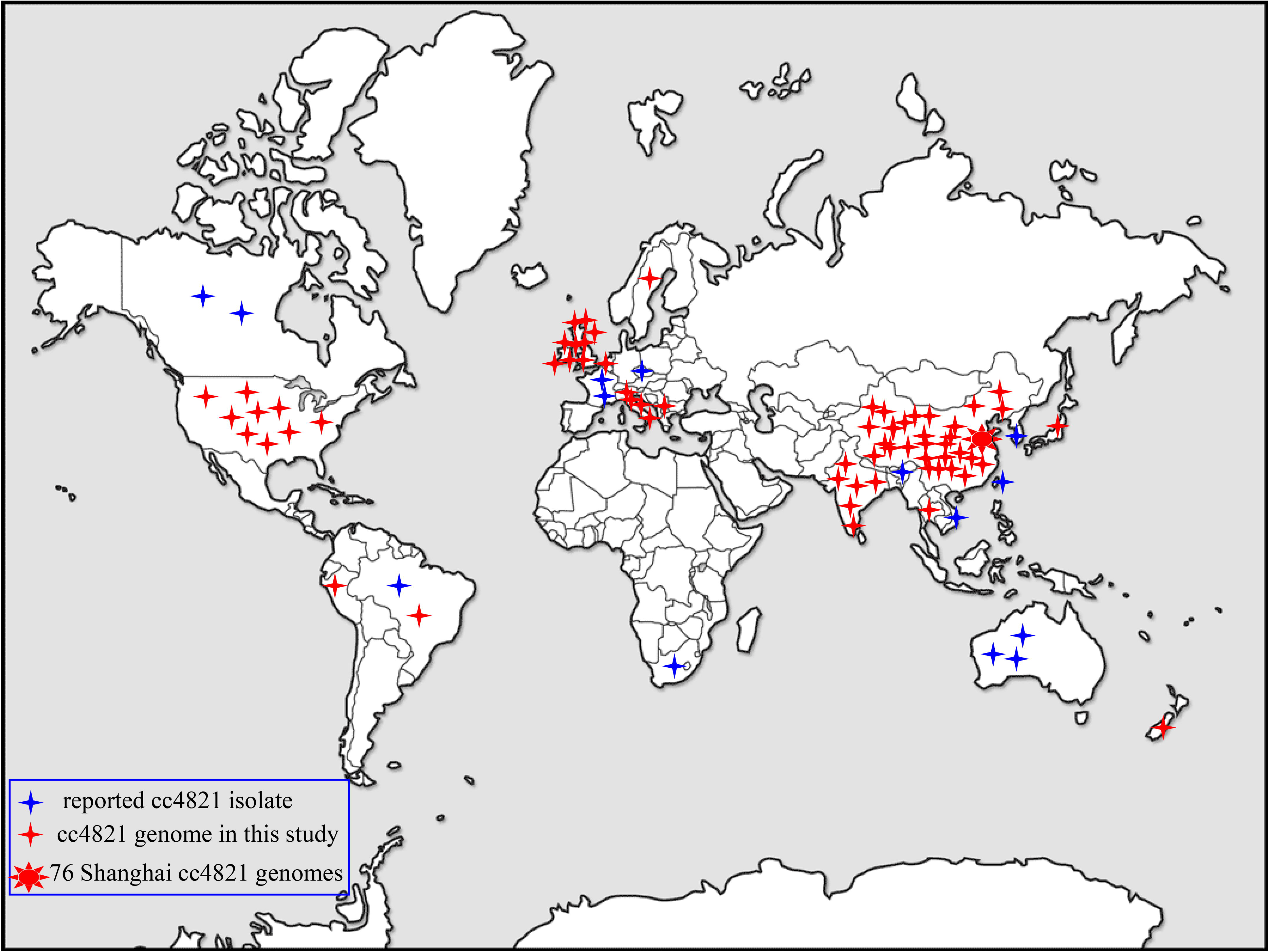
Distribution of ST-4821 clonal complex (cc4821) *Neisseria meningitidis* isolates worldwide. Besides China, cc4821 isolates were identified in other 19 countries of Europe, Africa, North America, South America, Oceania, and Asia.

## Materials and Methods

### Isolate collection and whole genome sequencing (WGS)

A total of 173 cc4821 genomes were collected for this study dating from 1972 to 1978 (n=19) and 2004 to 2019 (n=154), including isolate from IMD (37.6%, 65/173), non-invasive infections (4.0%, 7/173), asymptomatic carriage (48.0%, 83/173), and unknown sources (10.4%, 18/173) (Table S1). Shanghai CDC collected and sequenced 76 cc4821 isolates with Illumina HiSeq using paired-end 150 base reads as previously described (*13*). An additional 97 publicly available cc4821 genomes consisted of: (i) 48 genomes from a further 14 Chinese provinces, including the genome of MenC reference strain 053442 (*6, 8, 14*); (ii) 49 isolates from countries outside of China, including UK (n=20), USA (n=8), and 11 other countries (n=21) (Figure 1 and Table S1) (*10, 12, 15–18*). The completeness and contamination of the genomes were evaluated using CheckM (*19*).

### Antigenic and antibiotic resistance characteristics of the cc4821 genomes

To describe the antigenic and antibiotic resistance characterisations of cc4821 genomes, nucleotides of 9 antigen coding genes (*porA*, *fHbp*, *nhba*, *porB*, *fetA*, *opcA*, *nspA*, *tbpA*, and *NMB0315*) (*20–23*) and 5 antibiotic resistance-associated genes (*gyrA*, *parC*, *penA*, *ponA*, and *rpoB*) (*24–25*) were extracted from genomes for analysis. Deduced fHbp, NHBA, NadA, and PorA peptides and deduced meningococcal vaccine antigen reactivity (MenDeVAR) Index were annotated and analysed from PubMLST.org *Neisseria* website.

### Identification of the cc4821 (L44) sub-lineages

In *Neisseria* PubMLST database, a lineage-specific core genome MLST typing scheme containing loci found in 95% of cc4821 isolates was established and designated L44 cgMLST since cc4821 genomes were assigned to Lineage 44 (*26*). All of the 173 cc4821 genomes were compared using Genome Comparator and the L44 cgMLST scheme, allowing distinct sub-lineages to be identified. To identify features of each sub-lineage, a FASTA output from the Genome Comparator Tool using all the 2860 defined loci in the database (NEIS0001-NEIS3173, not contiguous) was visualized using MEGA v5 (*27*). The MenA meningococcus, Z2491 (Accession number NC_003116), was used as outgroup according to previous studies (*6, 8*).

### Identification and characterisation of unique alleles in sub-lineages

Outputs from Genome Comparator allowed shared and unique alleles to be determined among genomes. An allele was defined as unique to a sub-lineage if it was present in >90% of the genomes in that sub-lineage but absent in other sub-lineages. Genes with unique alleles were functionally characterised according to the Kyoto Encyclopedia of Genes and Genomes (KEGG) Orthology (KO) groupings of the KEGG database (*28*).

### Identification of HGT events and putative donors

Inputting the aligned sequences generated from Parsnp (*29*), putative HGT events were predicted using Gubbins (*30*). To search for potential donors, alleles and sequences of contiguous loci which were predicted as likely to originate from HGT were blasted against the PubMLST database. Potential donors for the unique alleles were identified as previously described (*31*). Recombination areas located with unique loci were labelled on the circular genome map of genome 053442 with BLAST comparisons with strains of other sub-lineages, generated using BRIG (*32*).

### Screening of the molecular markers of European MSM outbreak strains

Besides the lineage of 11.2 possessing PorA P1.5-1,10-8, three other molecular features were found in the meningococci causing outbreaks among MSM in Europe during 2012-14, comprising functional nitrite reductase (AniA), frameshifted fHbp allele, and *penA327* (with reduced susceptibility to penicillin and third-generation cephalosporins) (*33*). These three molecular markers were screened among all the 173 cc4821 genomes.

### Data access

Assembled contigs and annotation information of 173 genomes in this study can be accessed at https://pubMLST.org/neisseria using the IDs given in Table S1.

## Results

### Isolate characterisation

The 173 cc4821 isolates represented 46 different MLST sequence types (STs), with ST-4821 (n=41, 23.7%) and ST-3200 (n=30, 17.3%) the most prevalent. There were 43 PorA subtypes, of which P1.7-2,14 (n=25, 14.5%) and P1.17-6,23 (n=18, 10.4%) were the most frequent. There were 27 FetA variants, with F3-3 (n=47, 27.2%) and F3-36 (n=37, 21.4%) the most prevalent (Table S1).

### Identification of four sub-lineages

A total of 2161 loci were identified in reference genome 053442, including 1699 core genes. The majority (89.9%, 1527/1699) of the core loci had *p*-distance values of 0-0.1, and 0.8% (14/1699) showed high *p*-distance values of 0.50-0.68. Based on a lineage core genome scheme (L44 cgMLST) the cc4821 isolates were divided into 4 sub-lineages (Figure 2), termed (i) L44.1 (n=50, 28.9%; identical to the China^CC4821-R1-C/B^ clone), which comprised isolates from China (n=44) and other countries (n=6) during 2004-2019 and were very closely related; (ii) L44.2 (n=29, 16.8%), composed of isolates from China (n=28) and UK (n=1) during 2005-2019; (iii) L44.3 (n=58, 33.5%), comprising isolates from countries outside China (n=40) and China (n=18) during 1977-2019; (iv) L44.4 (n=32, 18.5%), composed of isolates from China (n=30) and India (n=2) during 1972-2017; and 4 other isolates from China not assigned to any sub-lineages.

**Figure. 2.**
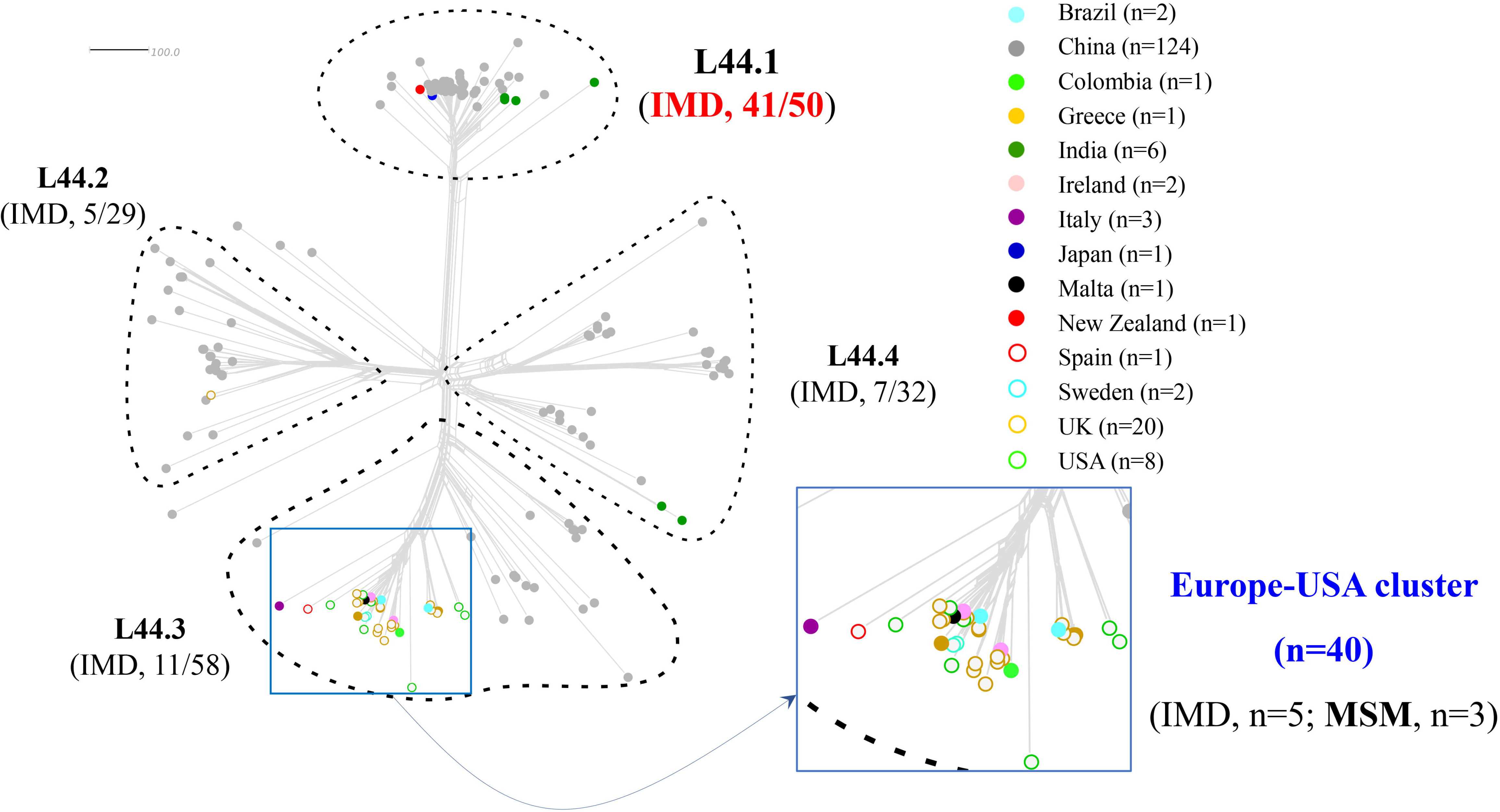
Allele-based sub-lineages of cc4821 clonal *N. meningitidis* identified using Lineage 44 core genome MLST scheme. IMD, invasive meningococcal disease; MSM, men who have sex with men.

### Features of the four different sub-lineages

The percentage of IMD isolates was significantly higher in L44.1 (82%, 41/50) than other three sub-lineages (17.2-22.4%, *P* < 0.001; Figure 2). L44.1, containing the reference strain 053442, was mainly composed of MenC isolates (88%, 44/50), with ST-4821 as the central ST; L44.2, MenB (93.1%, 27/29), ST-5664; L44.3, MenB (94.8%, 55/58), ST-3200; L44.4, MenC (43.8%, 14/32) and MenW (34.4%, 11/32), ST-3436 (Figure S1). Analysis of the 5 antibiotic resistance genes identified *gyrA*-71 (with T91I) and *parC*-12 were both unique to L44.1, *parC*-275 and *penA*-9 (with 5 mutations) were both unique to L44.3, while *gyrA*-294 (with T91I) was only discovered in L44.4 (Table 2, Figure S13-S17). In sub-lineage L44.1, all the isolates possessed the quinolone resistance-associated mutation T91I in GyrA (Figure 3). In sub-lineage L44.3, 69.0% (40/58) harboured PBP2 mutations, which were almost from countries outside of China (38/40, 95%) (Figure 4).

**Figure 3.**
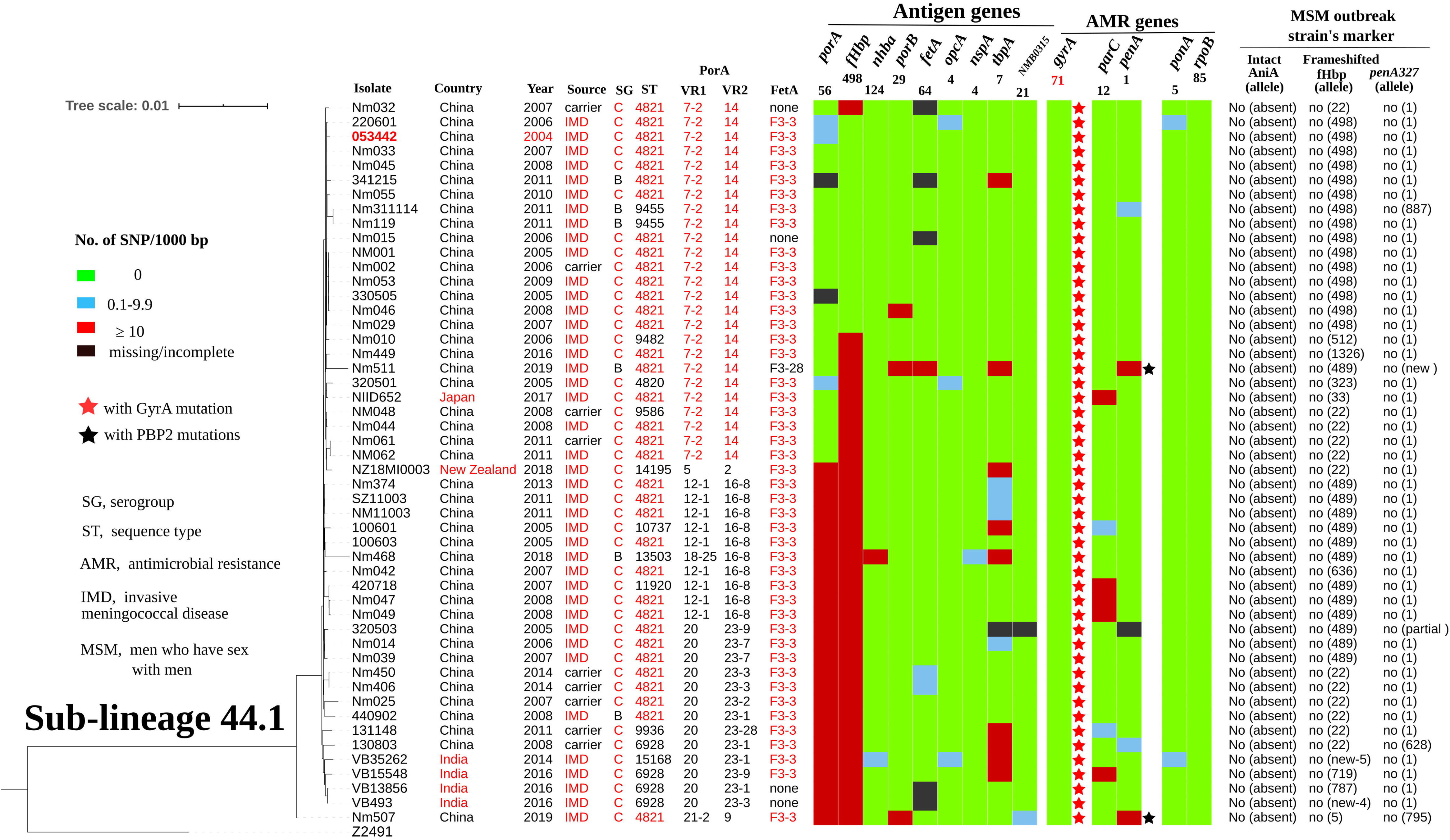
Genomic diversity of sub-lineages L44.1 (China^CC4821-R1-C/B^) isolates.

**Figure 4.**
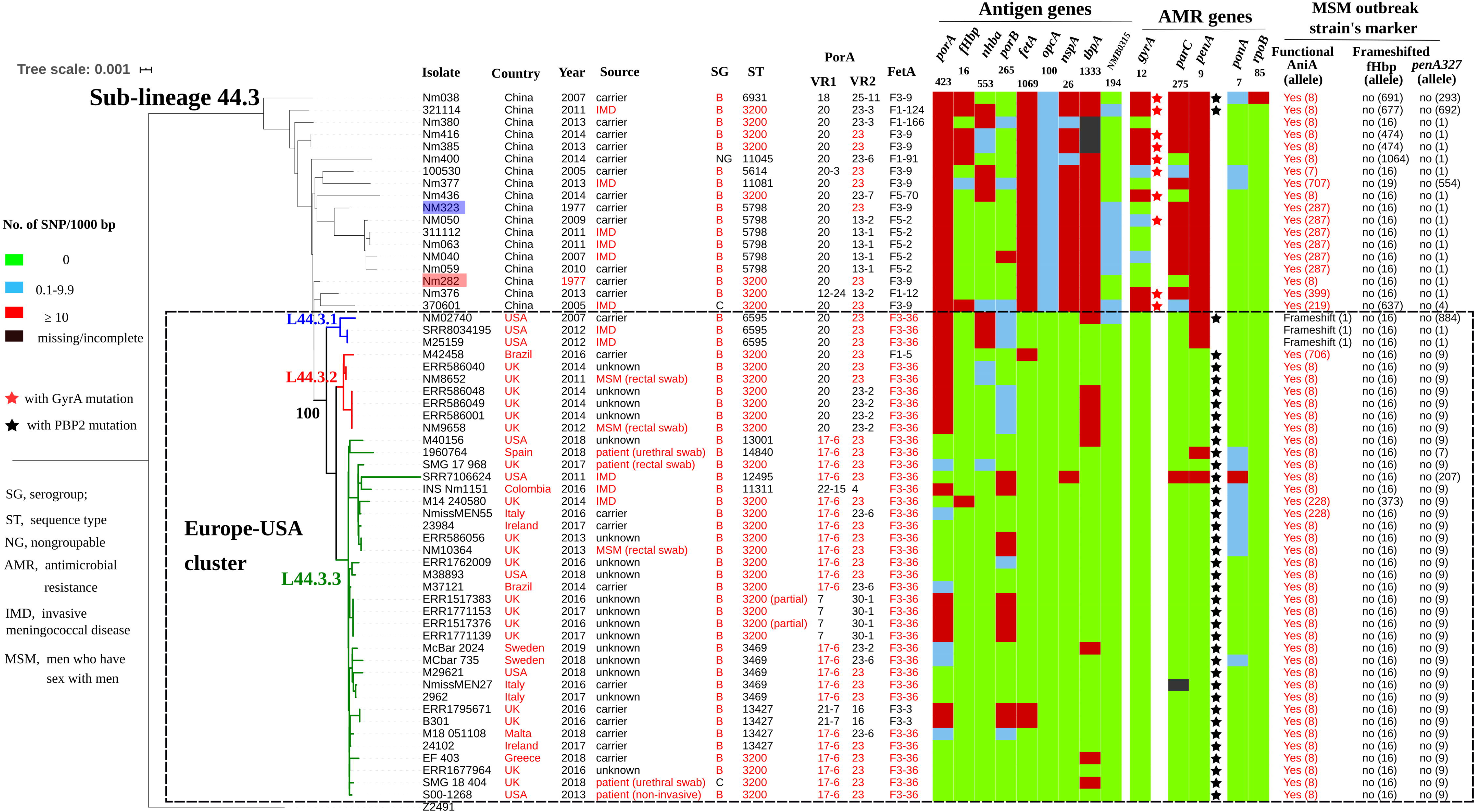
Genomic diversity of sub-lineages L44.3 isolates. The Europe-USA cluster can be further divided into three sub-clusters: sub-cluster L44.3.1 was composed of three USA ST-6595 isolates, all of which contained putatively non-functional AniA; L44.3.2 grouped seven ST-3200 isolates from UK (n=6) and Brazil (n=1); while L44.3.3 comprised 30 isolates with multiple geographic locations (Figure 4). All the isolates from urethral (n=2) and rectal (n=4) swabs were assigned to L44.3.2 and L44.3.3, both of which comprised isolates with putatively functional AniA.

### Vaccine antigens amongst the four sublineages

Analysis of the 9 antigenic genes identified several alleles that were unique to a certain sub-lineage (Table 2, Figure S2-S12). For example, FetA-VR F3-3 was only found in L44.1, F1-91 to L44.2, F3-36 and F3-9 in L44.3 isolates, and F1-7 in L44.4 (Figure S6). In L44.1, the majority had the same antigenic gene profile (*nhba*-124, *porB*-29, *fetA*-64, *opcA*-4, *nspA*-4, *tbpA*-7, and *NMB0315*-21, and 50% (25/50) had the PorA subtype of P1.7-2,14 (Figure 3). In sub-lineage L44.3 isolates, the majority had the same gene profile (*fHbp*-16, *nhba*-553, *porB*-265, *fetA*-1069, *opcA*-100, *nspA*-26, and *NMB0315*-335), with *porA* and *tbpA* showing high genetic diversity (Figure 4).

Deduced peptide sequences were analysed for vaccine antigen constituents amongst MenB isolates (n=97). There were 16 fHbp peptides, of which peptide 16 (variant 2/subfamily A) was present in 70/97 (72.2%) isolates, including 31/70 isolates from China. There were 20 NHBA peptides, of which 669 (48.4%, 46/95), 901 (11.6%, 11/95) and 668 (10.5%, 10/95) occurred most frequently. The *nadA* gene was absent in all isolates (including other serogroups). There were 31 unique PorA VR1/VR2 combinations, the most frequently occurring were P1.20,23 (11.3%, 11/97).

The MenDeVAR Index was assigned for MenB disease isolates (n=47), but 45/47 (95.7%) isolates had insufficient data from experimental studies to estimate if Bexsero^^®^^ would be reactive (Table S2). For Trumenba^®^, 35/47 (74.5%) isolates were predicted to have a cross reactivity based on experimental data from Meningococcal Antigen Surface Expression (MEASURE) assay and serum bactericidal activity assays (*34*). For the Chinese MenB disease isolates, 7/17 (41.2%) were deemed cross-reactive but the remaining 10/17 (58.8%) with insufficient data to determine reactivity.

### Screening of the molecular markers of European MSM outbreak strains

None of the cc4821 isolates harboured frameshifted fHbp allele or *penA327*, but the presence of functional AniA was diverse. The *aniA* gene was absent in all the L44.1 isolates, and was present in all the other 123 cc4821 isolates, of which only 96.7% (119/123) isolates harboured putatively functional AniA (Figure 3, 4, S2, and S3).

### Evolution of sub-lineage L44.1 (China^CC4821-R1-C/B^ clone)

Five specific loci were present in more than 90% of sub-lineage L44.1 while in less than 10% of other sub-lineages. These loci were involved in signaling and cellular processes (n=2), metabolism (n=1), and Genetic information processing (n=1) (Table 1). There were no loci specific to any of other three sub-lineages.

**Table 1.**
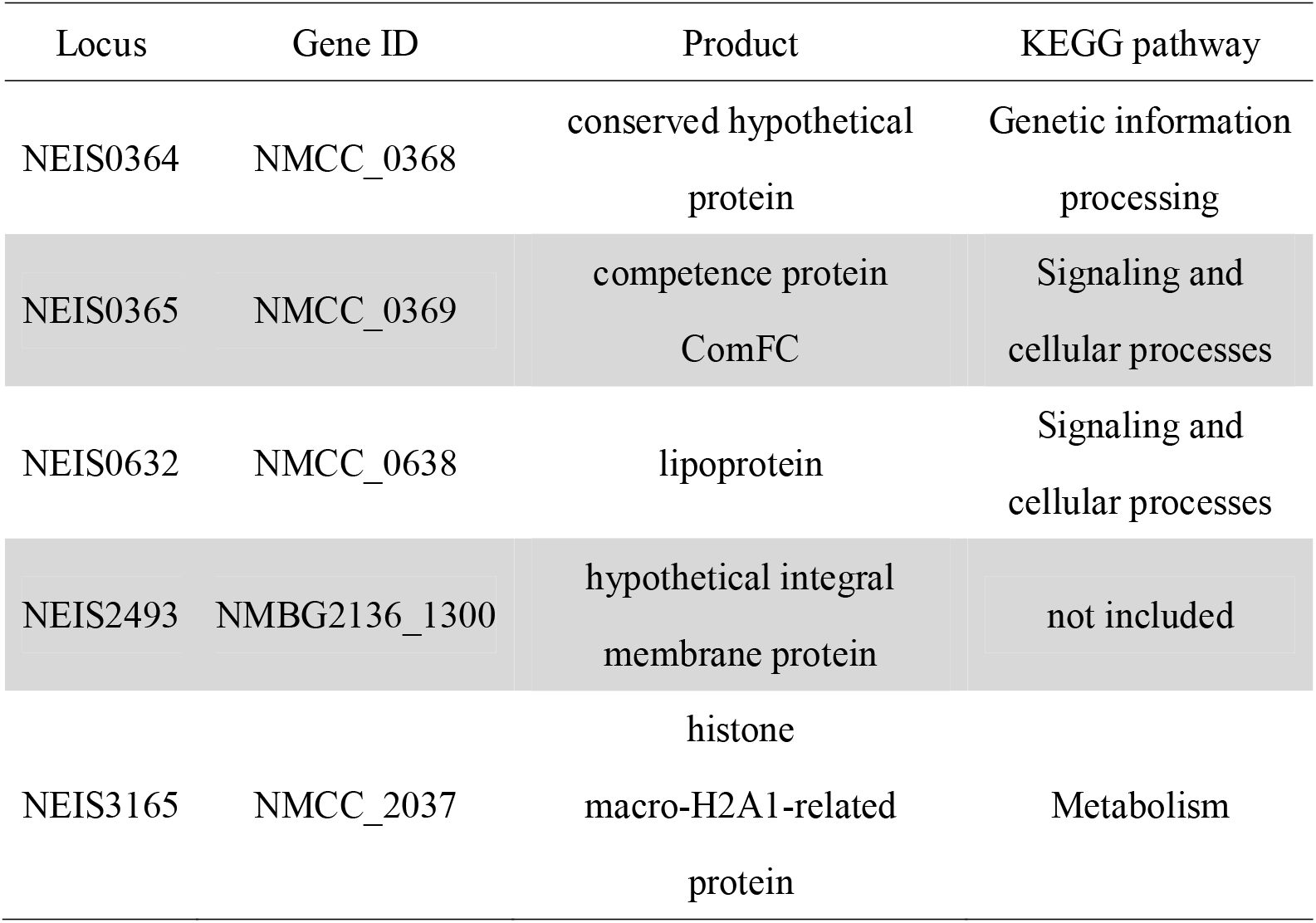
Loci only present in sub-lineage L44.1.

| Locus | Gene ID | Product | KEGG pathway |
| --- | --- | --- | --- |
| NEIS0364 | NMCC_0368 | conserved hypothetical protein | Genetic information processing |
| NEIS0365 | NMCC_0369 | competence protein ComFC | Signaling and cellular processes |
| NEIS0632 | NMCC_0638 | lipoprotein | Signaling and cellular processes |
| NEIS2493 | NMBG2136_1300 | hypothetical integral membrane protein | not included |
| NEIS3165 | NMCC_2037 | histone macro-H2A1-related protein | Metabolism |

**Table 2.**
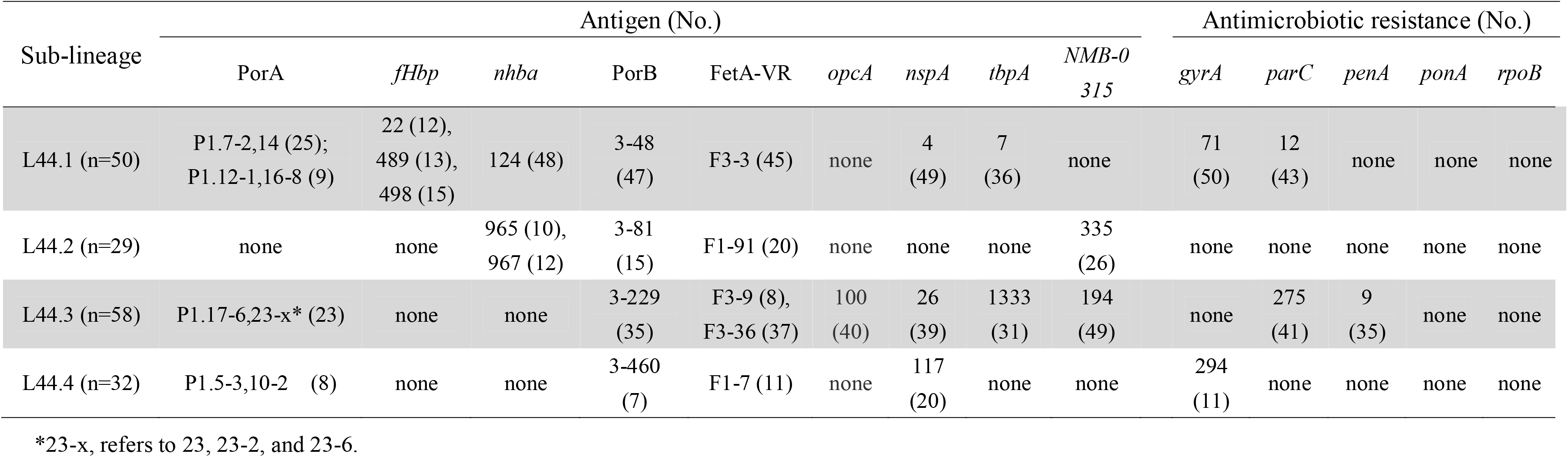
Specific alleles of antigenic and antimicrobial resistance genes in 4 sub-lineages of cc4821.

| Sub-lineage | Antigen (No.) |  |  |  |  |  |  |  |  | Antimicrobial resistance (No.) |  |  |  |  |
| --- | --- | --- | --- | --- | --- | --- | --- | --- | --- | --- | --- | --- | --- | --- |
|  | PorA | *fHbp* | *nhba* | PorB | FetA-VR | *opcA* | *nspA* | *tbpA* | *NMB-0315* | *gyrA* | *parC* | *penA* | *ponA* | *rpoB* |
| L44.1 (n=50) | P1.7-2,14 (25);<br>P1.12-1,16-8 (9) | 22 (12),<br>489 (13),<br>498 (15) | 124 (48) | 3-48<br>(47) | F3-3 (45) | none | 4<br>(49) | 7<br>(36) | none | 71<br>(50) | 12<br>(43) | none | none | none |
| L44.2 (n=29) | none | none | 965 (10),<br>967 (12) | 3-81<br>(15) | F1-91 (20) | none | none | none | 335<br>(26) | none | none | none | none | none |
| L44.3 (n=58) | P1.17-6,23-x* (23) | none | none | 3-229<br>(35) | F3-9 (8),<br>F3-36 (37) | 100<br>(40) | 26<br>(39) | 1333<br>(31) | 194<br>(49) | none | 275<br>(41) | 9<br>(35) | none | none |
| L44.4 (n=32) | P1.5-3,10-2 (8) | none | none | 3-460<br>(7) | F1-7 (11) | none | 117<br>(20) | none | none | 294<br>(11) | none | none | none | none |

\*23-x, refers to 23, 23-2, and 23-6.

In prediction of HGT events contributing to the emergence of sub-lineage 44.1 by Gubbins, 126 events involving 686 loci were discovered to be shared by the 50 L44.1 isolates (Figure S18), including 216 loci with alleles specific (unique loci) to L44.1. Another 83 unique loci were discovered based on analysis of accessory loci. Therefore, a total of 299 unique loci were identified in L44.1, 46.5% (139/299) of which were involved in metabolic function (Table S3).

These 299 unique loci were distributed across the chromosome, and a total of 44 areas (216 loci) harbouring contiguous loci with unique alleles were observed (Figure 5), among which the exact donors of 36 areas across 149 loci were identified in 46 putative HGT events. The total length of these putative recombination fragments was approximately 225 kb, including 87 kb (38.7%) originating from the C:ST-9514 cluster isolates in China during 1966-1977, followed by 25 kb (11.1%) from MenA isolates (cc5 and cc1) in China during 1966-1984 (Table 3).

**Figure 5.**
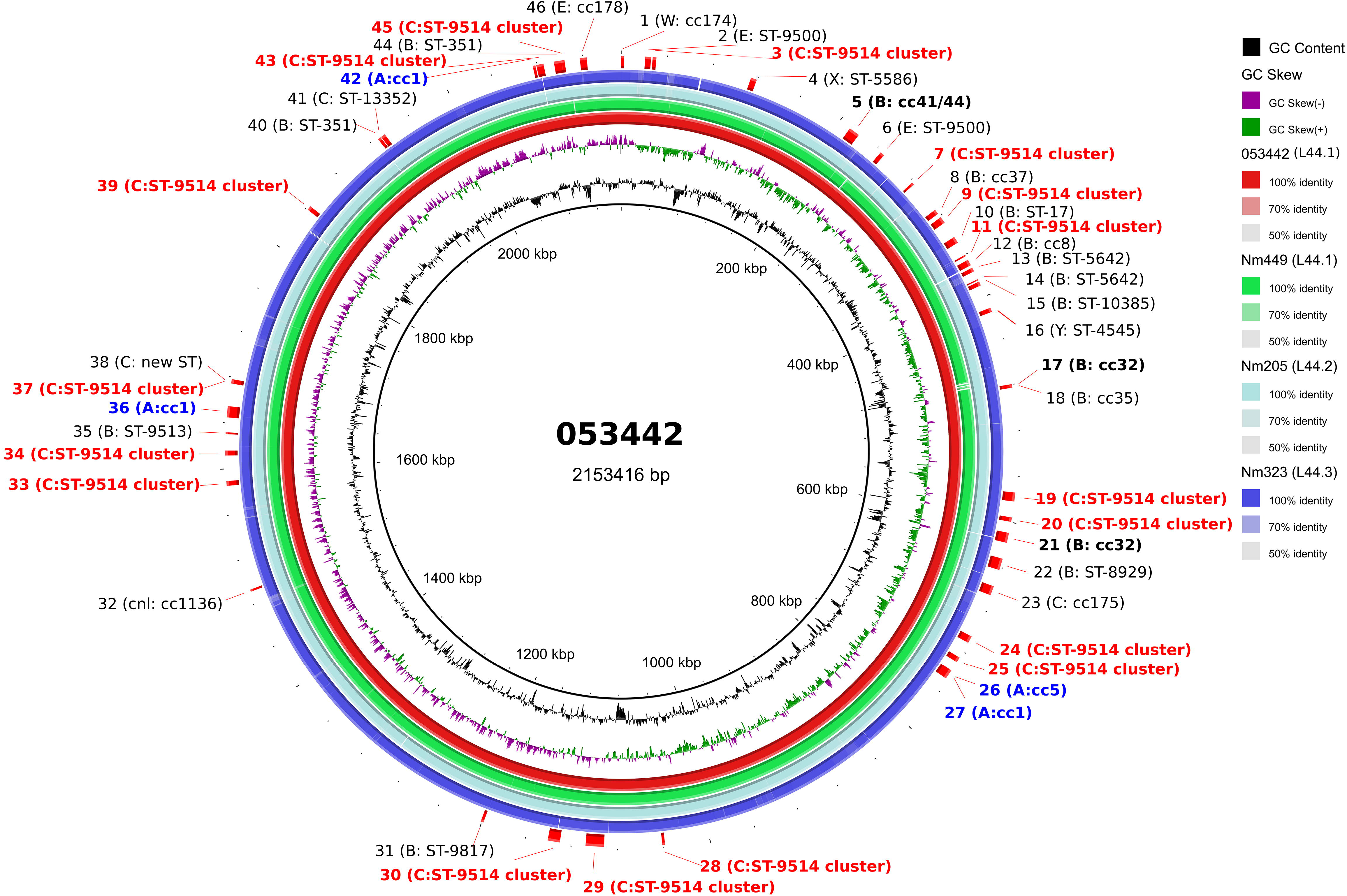
Circular genome map of cc4821 *N. meningitidis* genome 053442 with BLAST comparisons to the genomes of other sub-lineages. The innermost rings show G+C content (black) and G+C skew (purple/green) of 053442. The 4 outer rings show BLAST comparisons (using BLASTn and an E-value cutoff of 10.0) to the complete genome sequence of 053442 (red), Nm449 (green), Nm205 (pale blue), and Nm323 (blue). Labels around the outer ring refer to the 46 HGT events involving 149 unique loci which are labeled with their possible donor strain.

**Table 3.**
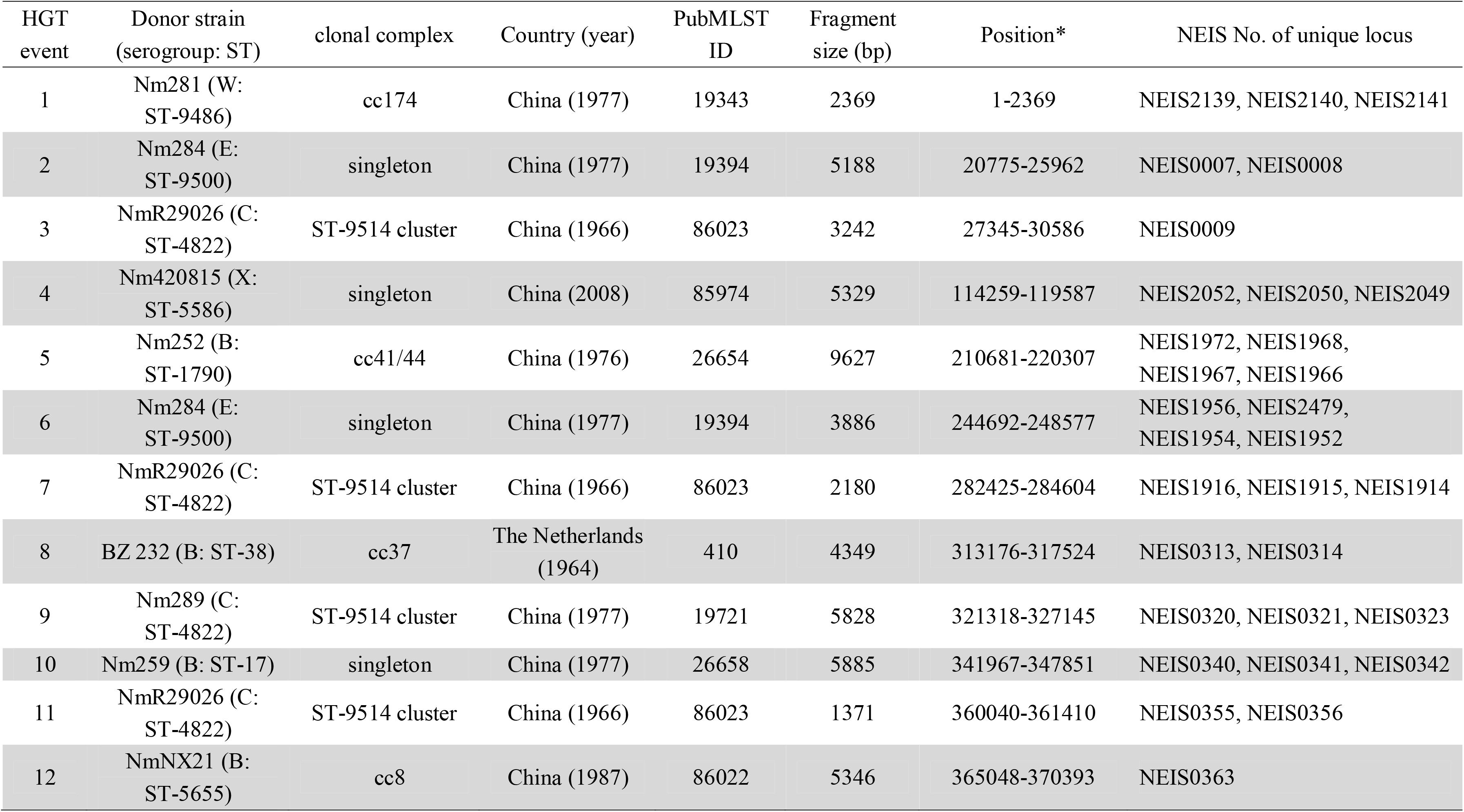
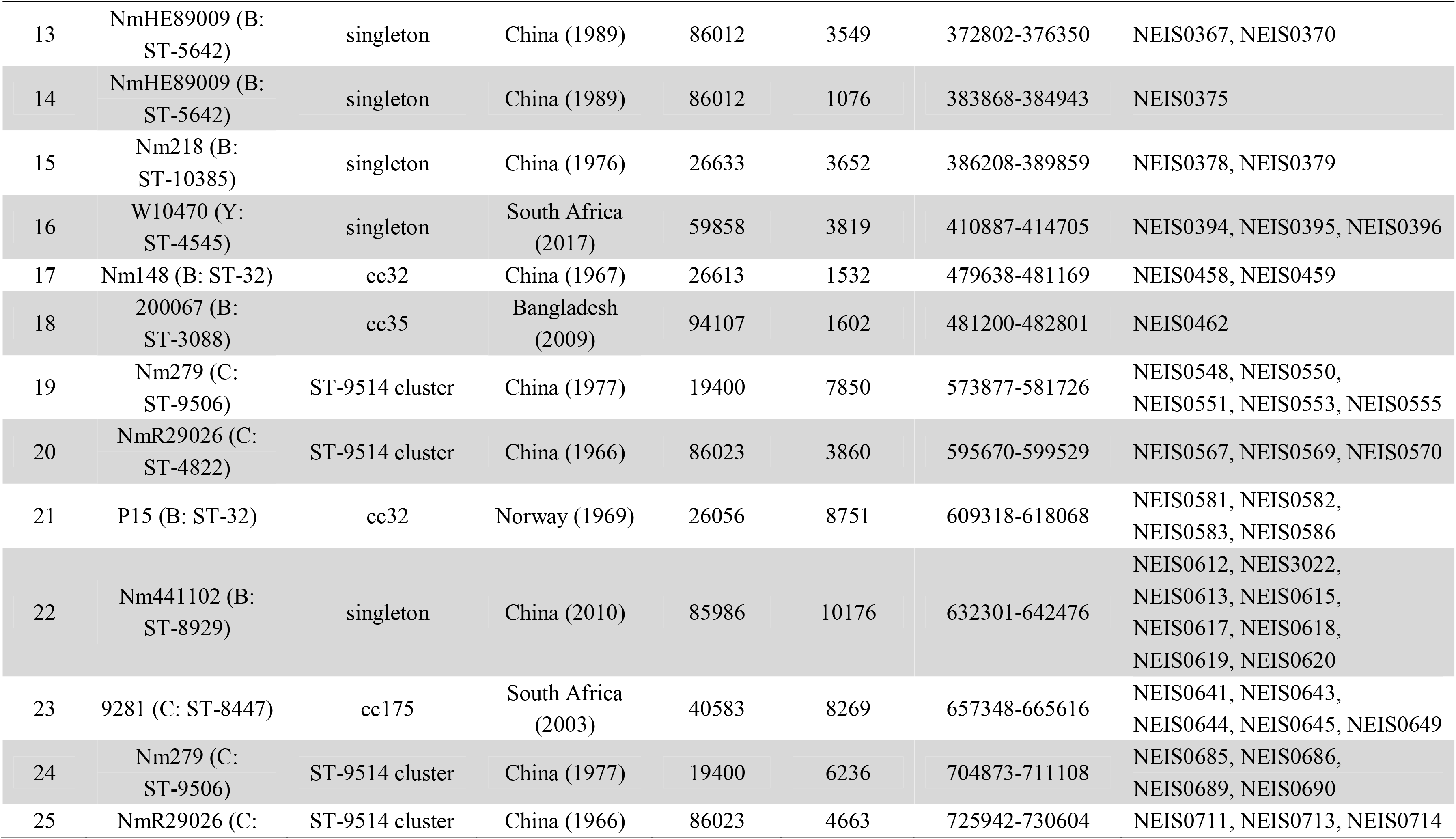
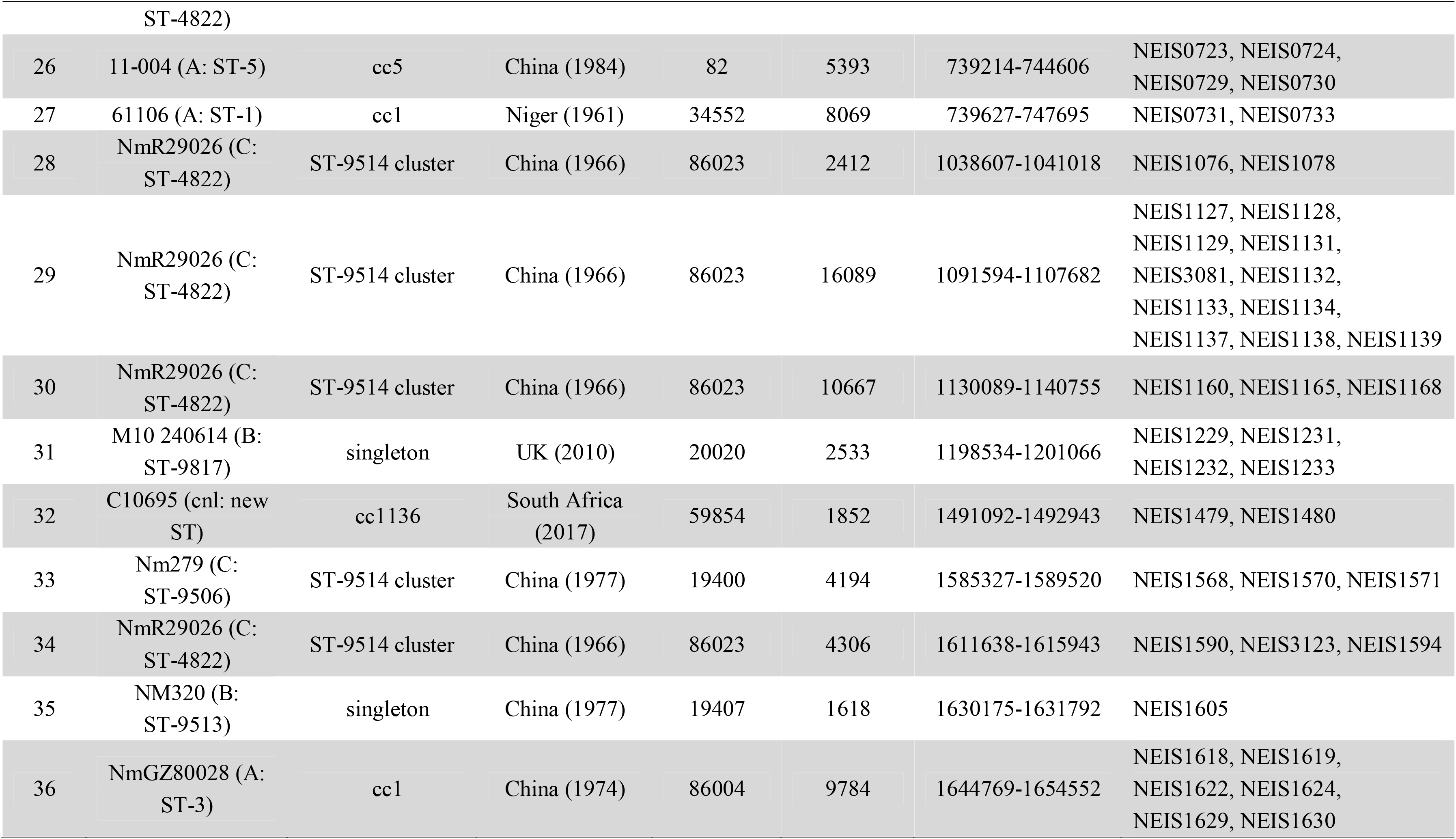
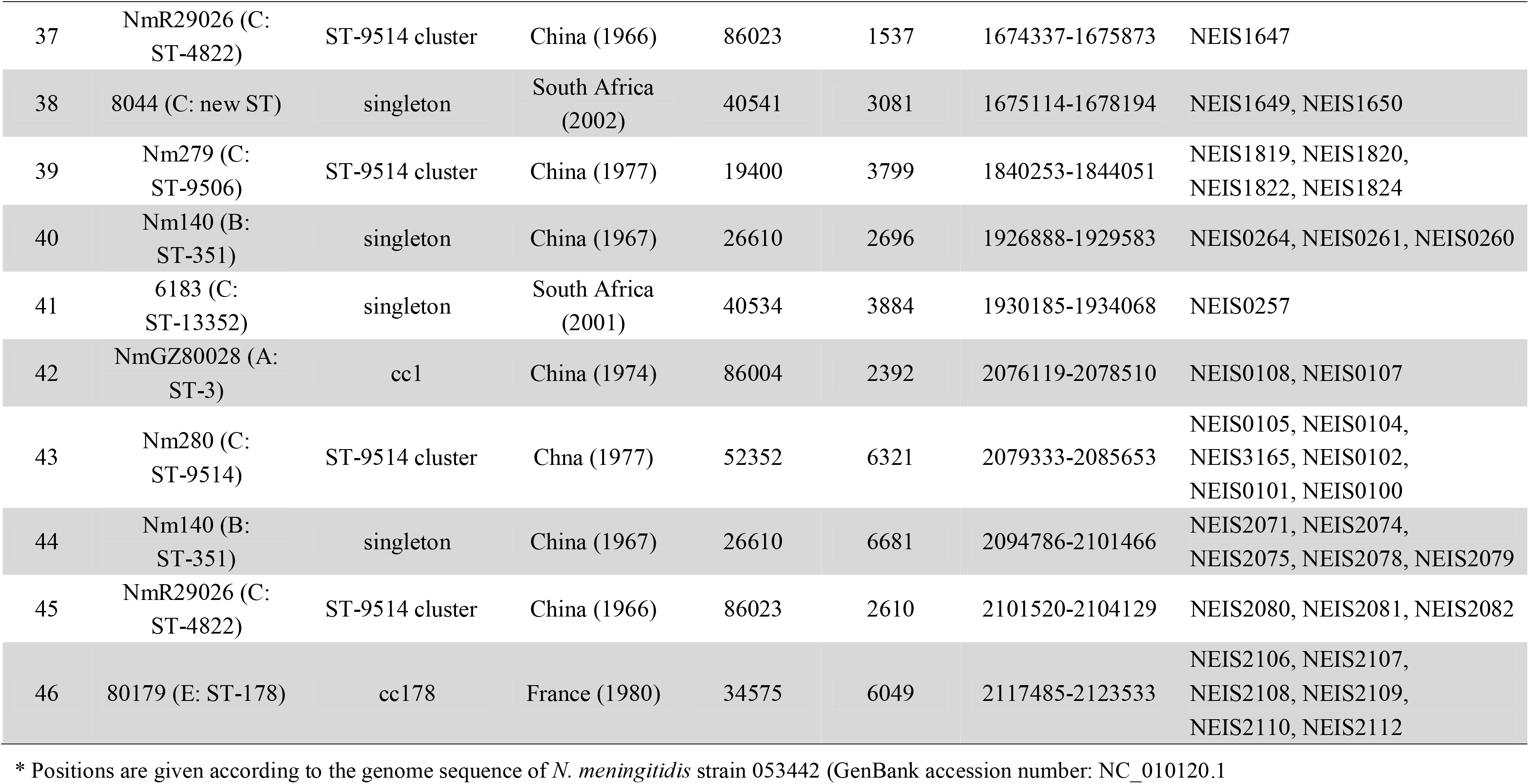
Potential donors of horizontal gene transfer (HGT) events associated with contiguous unique loci in sub-lineage 44.1.

### Evolution of cc4821 isolates from other countries

There were 49 cc4821 isolates from countries outside of China, and most (81.6%, 40/49) were assigned to L44.3, in which these 40 isolates, including 39 MenB and 1 MenC, constituted a distinct cluster (the Europe-USA cluster, Figure 2 and 4). The representative molecular characteristics of the Europe-USA cluster was B:P1.17-6,23: F3-36:ST-3200(cc4821), with the gene profiles of antigen (*porA*-423, *fHbp*-16, *nhba*-553, *porB*-265, *fetA*-1069, *opcA*-100, *nspA*-26, *tbpA*-1333, and *NMB0315*-194) and antibiotic resistance (*gyrA*-12, *parC*-275, *penA*-9 with mutations in PBP2, *ponA*-7, and *rpoB*-85) (Figure 4). In Gubbins analysis, 33 events involving 193 loci were shared by all the Europe-USA cluster isolates (Figure S4), and 60 unique loci were discovered but without clues to their potential donors. These unique loci were involved in functions mainly of metabolism (38.3%, 23/60) and genetic information processing (30%, 18/60) (Table S4).

Except the 40 Europe-USA cluster isolates, six MenC invasive isolates from other countries, including India (n=4, 2014-2016), Japan (n=1, 2017), and New Zealand (n=1, 2018), were clustered together and closely related with 44 Chinese isolates within the same sub-lineage, L44.1 (Figure 2 and 4). Only the Japanese isolate showed the typical molecular feature of Anhui outbreak strain (C:P1.7-2,14:F3-3:ST-4821[cc4821]).

### Features and evolution of serogroup W cc4821 isolates

There were 11 MenW isolates from China and the representative strain designation was W:P1.5-3,10-2:F1-7:ST-8491(cc4821), with similar gene profiles of antigen-encoding loci (*porA*-1804, *fHbp*-641, *nhba*-966, *fetA*-37, *opcA*-4, *nspA*-117, and *NMB0315*-21) and antibiotic resistance loci (*gyrA*-294 with T91I, *parC*-779, *ponA*-7, and *rpoB*-85). These MenW isolates constituted a distinct cluster in the L44.4, and they were closer to NM193 (C:P1.20-3,23-1:F1-5:ST-3436[cc4821], in 1972) than to NM205 (C:P1.20,23-2:F5-135:ST-4821[cc4821], in 1973;) (Figure S12).

## Discussion

Although primarily a member of the commensal microbiota, the meningococcus can cause IMD, leading to endemic disease in most if not all human populations. A number of genotypes, members of the hyperinvasive lineages, in combination with the disease-associated capsular serogroups can cause elevated levels of disease and some of these also have epidemic and pandemic potential. Over the past 100 years notable epidemic/pandemic meningococci have included: A:cc1; A:cc5; B:cc41/44; C:cc11; and W:cc11. Here we have employed genomic analysis of serogroup B, C, and W cc4821 meningococci isolated from 1972 to 2019 to assess their epidemic and pandemic potential. Of especial concern is: the expansion of the quinolone-resistant clone China^CC4821-R1-C/B^ from China to other countries; the potential possession of universal resistance to penicillin in Europe-USA cluster isolates; and the uncertainty over the potential efficacy of existing vaccines to prevent B:cc4821 IMD.

The cc4821 (corresponding to Lineage 44) shares several properties in common with the hyperinvasive cc11 (L11), including its: ability to express several serogroups; global distribution; colonisation in urogenital and anorectal tracts, and separation into distinct sub-lineages. cc11 has caused well-documented epidemics and pandemics on several occasions, including: the US military outbreaks in 1960s; the Hajj-associated outbreaks in 2000s; and the global epidemics from 2010 onwards, especially outbreaks among MSM (*33, 35–37*). These similar characteristics raise the concern that the cc4821 may have potential to cause global pandemics.

Consistent with the presence of the epidemic cc4821 clone in countries outside of China, four Indian, one Japanese, and one New Zealand cc4821 IMD meningococci, isolated during 2014-2018, were clustered with China^CC4821-R1-C/B^ meningococci in sub-lineage L44.1 (Figure 3). IMD cases caused by these six isolates were all in native inhabitants (*10, 16, 38*), and all six isolates shared similar characteristics of serogroup, antigen (except porA and fHbp), and antibiotic gene profile with Chinese isolates in this sub-lineage. Particularly, they all harboured T91I mutation in GyrA, the molecular marker of quinolone resistance, compatible with their quinolone-resistant phenotype (*10, 16, 38*). These six isolates also had strain-specific features, which suggests their being a result of transmission from the China^CC4821-R1-C/B^ clone. With the typical molecular features of the Anhui outbreak strain (C:P1.7-2,14:F3-3:ST-4821[cc4821]) (4), the Japanese isolate became the first reported quinolone-resistant meningococcus harbouring ParC mutation (S87I, allele 1538) to cause IMD worldwide (*10*). In contrast, none of the Chinese China^CC4821-R1-C/B^ clone had ParC mutations. Compared with the Japanese isolate, the four Indian and one New Zealand isolates were more distantly related to the reference strain 053442 (Figure 3), with different STs, *porA*, *fHbp,* and *tbpA* alleles.

Although a putative ancestor of the quinolone-resistant clone China^CC4821-R1-C/B^ was not identified in this study, we found 299 loci with alleles unique to this sub-lineage. Approximately half of these loci were associated with metabolic pathways, suggesting that divergence in metabolic genes may play an important role in the emergence of epidemic meningococci. Several studies have indicated that metabolic genes can influence the pathogenesis and virulence of the meningococcus, for example by allowing alternative host resources to be exploited in invasive disseminated infections (*39*). Changes in the hyperinvasive A:cc5 meningococci circulating in Africa have been associated with HGT of core genes involved in metabolic processes (*40*). The putative donors of these unique alleles included lineages from different serogroups and dates of isolation, such as: C:ST-9514 cluster, 1960s-1970s; A:cc5 and A:cc1, 1960s-1980s; B:cc32, 1960s; B:cc41/44, 1970s; and E:cc178, 1980s (Table 3). The C:ST-9514 cluster, which was defined previously as STs sharing at least 4 loci with ST-9514, was predominant in MenC carriage isolates during 1965-1980 in Shanghai, China (*41*). Therefore the emergence of China^CC4821-R1-C/B^ clone was perhaps associated with the accumulation of these unique alleles, which accounted for the separation from other sub-lineages in the allele-based phylogeny (Figure 2).

In the PubMLST database, over 60% of the cc4821 isolates from outside China were MenB. From these, 49 genomes were available in this study, including isolates from IMD (n=13) and urogenital and rectal tracts (n=6). The majority of these genomes clustered in the sub-lineage L44.3 and constituted a distinct cluster, the Europe-USA cluster, showing the typical strain designation: B:P1.17-6,23-x:F3-36:ST-3200(cc4821). The PorA and FetA types P1.17-6,23 and F3-36 were only found in this cluster. In the *Neisseria* PubMLST database, a further 24 cc4821 isolates from other countries (USA, Brazil, France, Czech Republic, Spain, Italy, Australia, and Vietnam) had no genome data, but with PorA or FetA variants (Table S5). Of these, 19 (79.2%) exhibited P1.17-6,23-x or F3-36, suggesting they might belong to the Europe-USA cluster. This cluster was distinct from the epidemic clone China^CC4821-R1-C/B^. For example, the antigen gene profile characteristic of the Europe-USA cluster was P1.17-6,23-x, F3-36, PorB-3-229, *fHbp*-16, *nhba*-553, *opcA*-100, *nspA*-26, and *tbpA*-1333, compared to P1.7-2,14, F3-3, PorB-3-48, *fHbp*-498, 22 and 489, *nhba*-124, *opcA*-4, *nspA*-4, and *tbpA*-7, in the China^CC4821-R1-C/B^ clone. In addition, all the China^CC4821-R1-C/B^ isolates harboured the mutation T91I in GyrA, while almost all of the Europe-USA cluster isolates possessed mutations in PBP2 (F504L, A510V, I515V, H541N, and I566V). This reflected different antibiotic selective pressures experienced by the Europe-USA and the China^CC4821-R1-C/B^ meningococci. A high frequency (>70%) of quinolone resistance has been reported in China since 2005 while 65% of meningococci in Europe showed reduced susceptibility to penicillin G during 1945-2006 (*5, 42–43*). In the two oldest isolates of the sub-lineage L44.3, Nm282 (B:P1.20,23:F3-36:ST-3200[cc4812]) was much closer to the Europe-USA cluster isolates than Nm323 (B:P1.20,23:F3-36:ST-5798[cc4821]) (Figure 4), and it seemed more likely to be the ancestor of the Europe-USA cluster isolates.

Urogenital and rectal meningococci have raised increasing public health concerns (*33*). In 2017, cc4821 anorectal isolates were identified in the UK (*12*). Here we identified cc4821 isolates from urethral or rectal tracts were clustered with those from IMD and oropharyngeal carriage (Figure 4), and most of cc4821 isolates contained putatively functional nitrite reductase (AniA) to grow in anaerobic environments, with the exception of L44.1 isolates. The cc4821 isolates acquired quinolone resistance alleles from *N. lactamica* and *N. subflava* (*44*), and the ability to grow in anaerobic environments will facilitate them to acquire gonococcal alleles, including antimicrobial resistance alleles. Such events seem to have already occurred to a sub-lineage of cc11, which was responsible for a number of IMD outbreaks and urethritis among MSM (*33*). They shared the same *penA* allele (*penA327/penAXXXIV*) with gonococci and showed decreased susceptibility to third-generation cephalosporins (*45*).

Lineage 44 (cc4821) includes isolates from different serogroups, including B, C, and W. In China, MenC and MenW isolates can be prevented by vaccines, such as group A and C meningococcal polysaccharide vaccine (MPV-AC) and MPV-ACYW, but there is no routinely administered vaccine to prevent MenB IMD (*46*). At present, two protein-based MenB vaccines, 4CMenB (Bexsero^^®^^) and rLP2086 (Trumenba^®^), have been licensed in several countries (*47–49*), but there are limited data on the bacterial coverage of these vaccines to cc4821 isolates directly from serum bactericidal activity assays or the MATS for Bexsero^®^ or MEASURE for Trumenba^®^ assay. Only one B:cc4821 isolate (M14-240580, UK) was reported to be tested using the MATS assay and showed no potential protection (*50*). Using systems to index complex genotypic and phenotypic data, for example the MenDeVAR Index, it was predicted that ~75% of B:cc4821 disease-causing isolates might be prevented through vaccination with Trumenba^^®^^, but there is insufficient data to infer Bexsero^^®^^ reactivity. Further testing of globally diverse meningococci is needed with these experimental assays to analyse potential vaccine impact in non-European settings.

In summary, a comprehensive genomic analysis of a hyperinvasive meningococcal cc4821 expressing serogroups B, C, and W with expansion from China to other global geographic locations has been undertaken. This identified key genomic factors and putative evolutionary changes, which might have led to the emergence and persistence of the epidemic quinolone-resistant clone in China. The vaccine coverage to the B:cc4821 isolates need further evaluation. Enhanced laboratory surveillance for cc4821 isolates from IMD cases, oropharyngeal, urethral and rectal carriage are needed to monitor its global trends of expansion, which will be essential for local immunisation policies.

## Acknowledgements

This study made use of Neisseria genomic data deposited in the *Neisseria* MLST Database (https://pubmlst.org/neisseria/) sited at the University of Oxford (Jolley & Maiden 2018, Wellcome open research, 3:124). The development of this database has been funded by the Wellcome Trust and European Union.

## Biographical sketch

Dr. Chen is an associate professor in Department of Microbiology, Shanghai Municipal Center for Disease Control and Prevention. His research interests include mechanisms of antimicrobial resistance in clinical isolates responsible for respiratory tract infections.

