## Supplementary figures for "Evolution of ST-4821 clonal complex hyperinvasive and quinolone-resistant meningococci: the next meningococcal pandemic?"

Technical Appendix

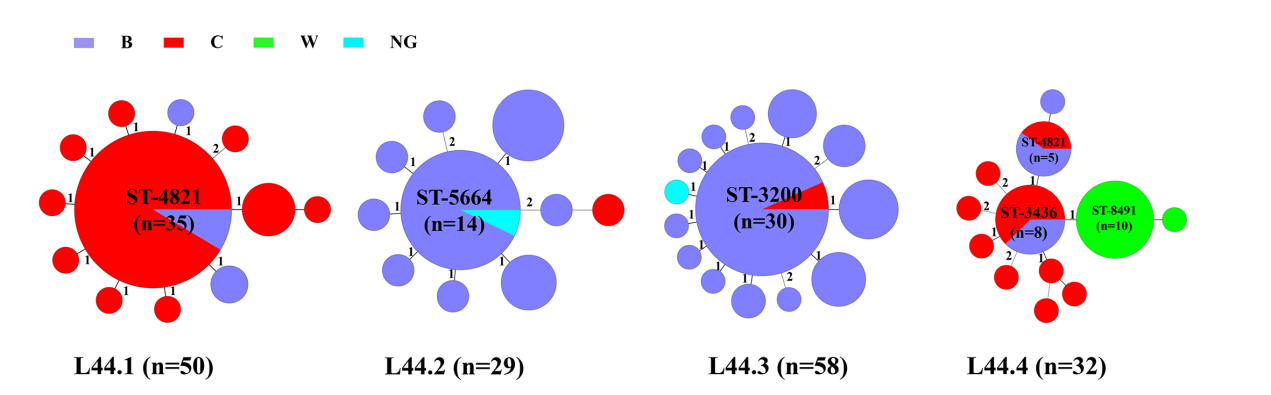

**Figure S1. Central sequence type of each sub-lineage of ST-4821 complex.**

The numbers of different loci between the central sequence types (ST) and its neighboring STs are labelled. The ST which differs only one locus to the central ST is a single-locus variant (SLV) of the central ST. In sub-lineage 44.1 (L44.1), ST-4821 (n=35) and its SLVs (n=13) take up for 96% (48/50); in L44.2, ST-5664 (n=14) and its SLVs (n=12) take up for 89.7% (26/29); in L44.3, the percentage of ST-3200 (n=30) and its SLVs (n=23) is 91.4% (53/58); in L44.4, the percentage of ST-3436 (n=8) its SLVs (n=18) is 81.3% (26/32).

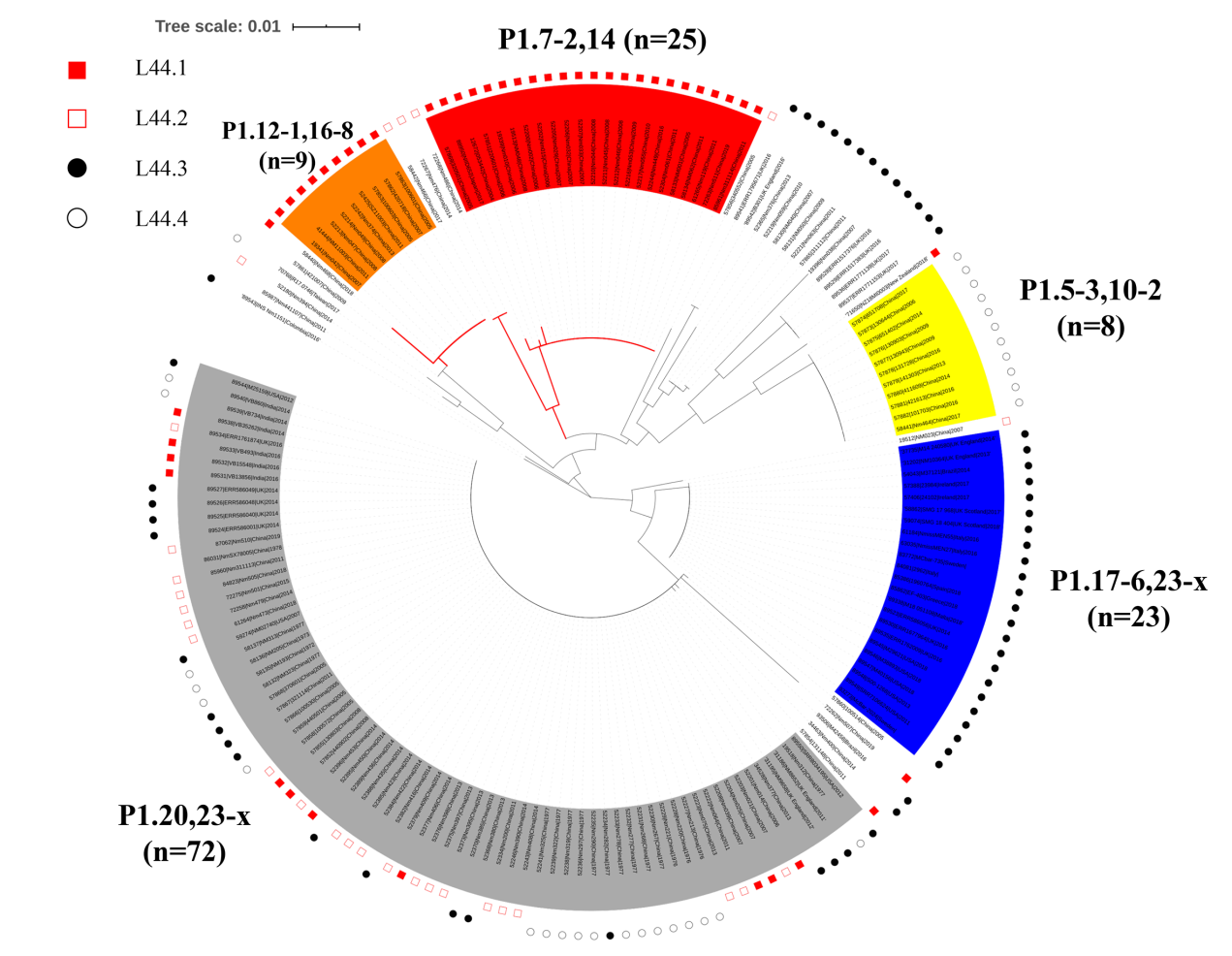

**Figure S2. Phylogenetic analysis of antigen gene sequence variation within the *N. meningitidis* clonal complex 4821 lineage, based on nucleotide sequences of the *porA* gene.**

23-x, refers to 23, 23-1, 23-2, 23-3, 23-6, 23-7, 23-9, 23-18, 23-19, and 23-28.

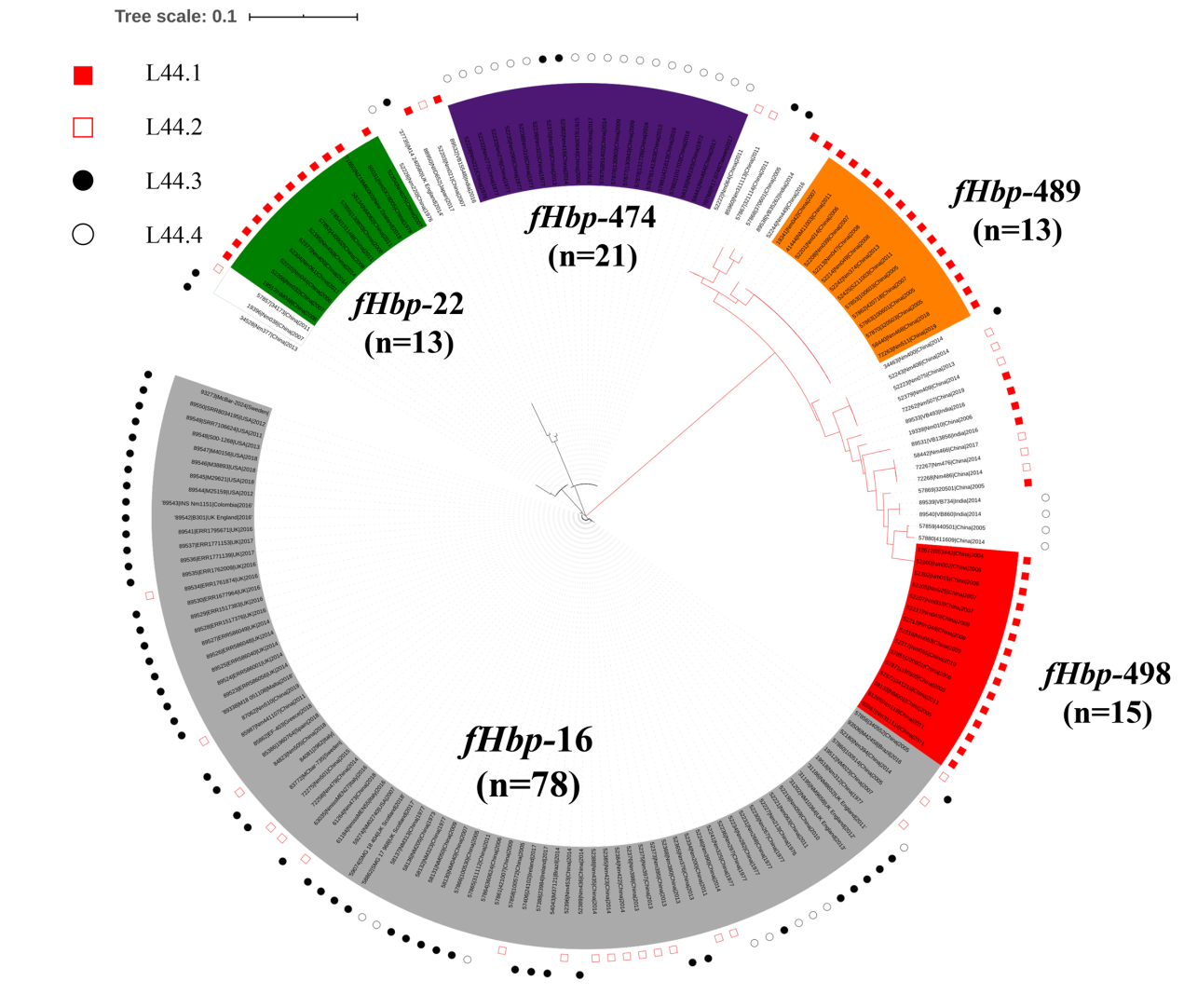

**Figure S3. Phylogenetic analysis of antigen gene sequence variation within the *N. meningitidis* clonal complex 4821 lineage, based on nucleotide sequences of the *fHbp* gene.**

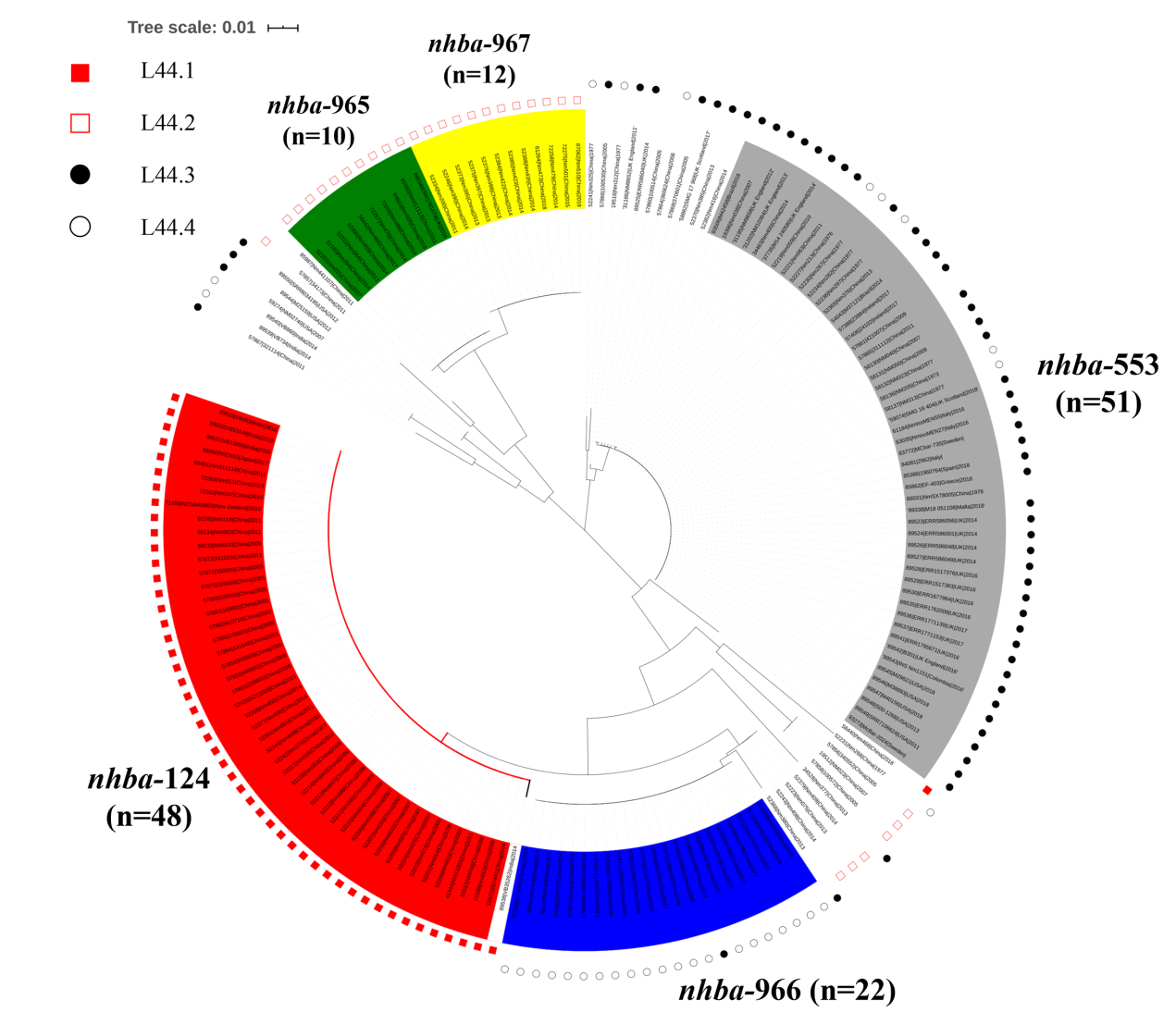

**Figure S4. Phylogenetic analysis of antigen gene sequence variation within the *N. meningitidis* clonal complex 4821 lineage, based on nucleotide sequences of the *nhba* gene.**

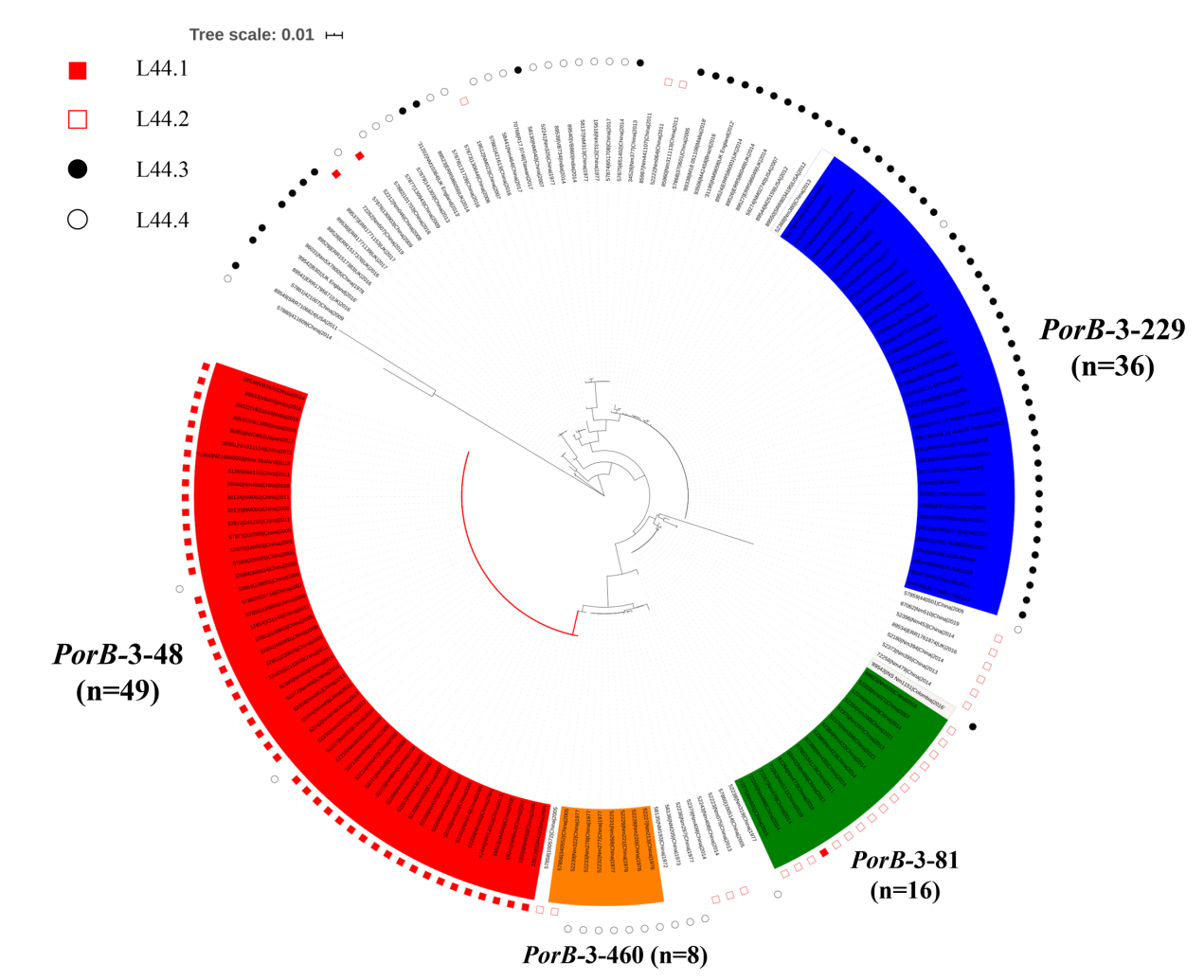

**Figure S5. Phylogenetic analysis of antigen gene sequence variation within the *N. meningitidis* clonal complex 4821 lineage, based on nucleotide sequences of the *porB* gene.**

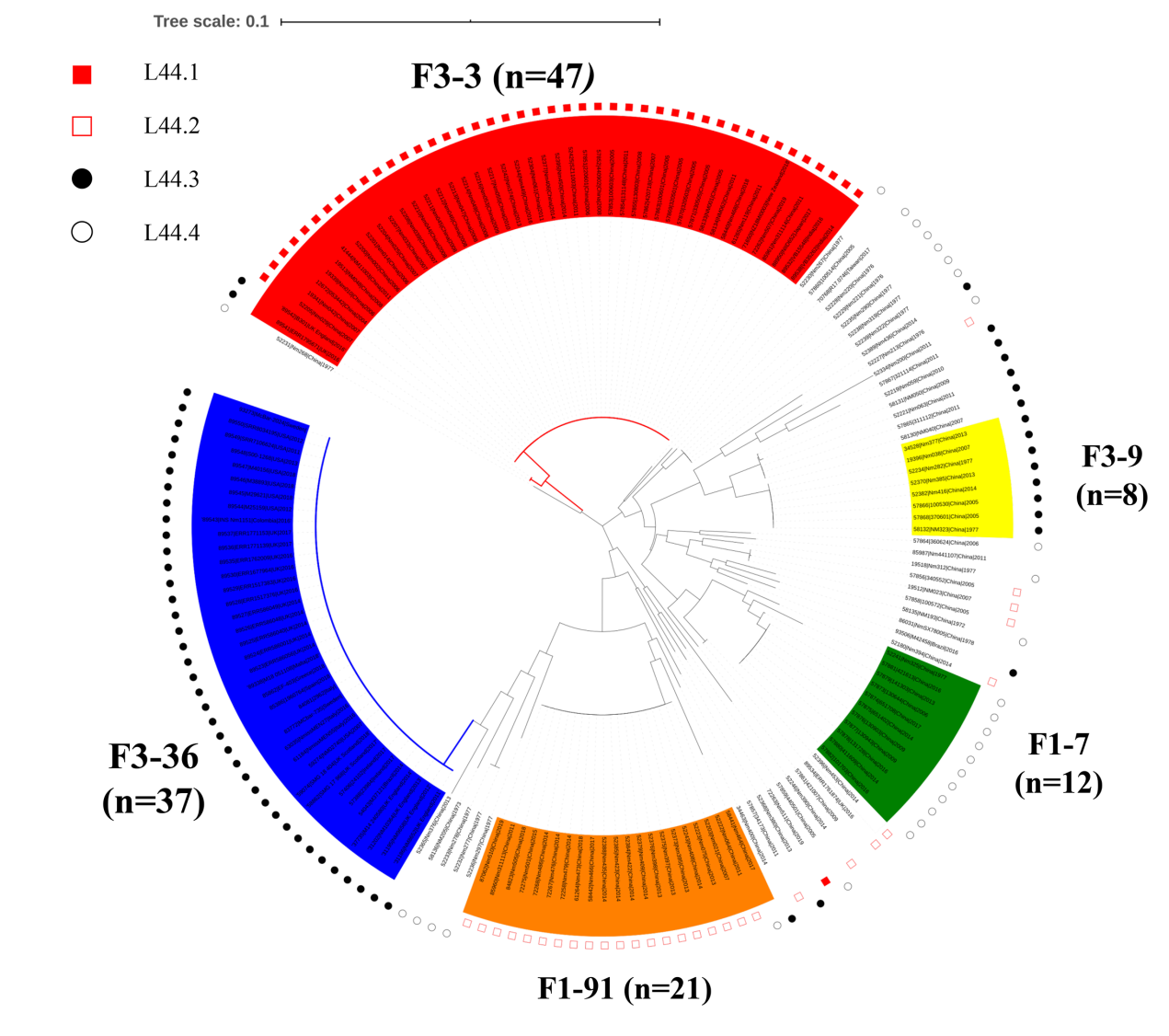

**Figure S6. Phylogenetic analysis of antigen gene sequence variation within the *N. meningitidis* clonal complex 4821 lineage, based on nucleotide sequences of the *fetA* gene.**

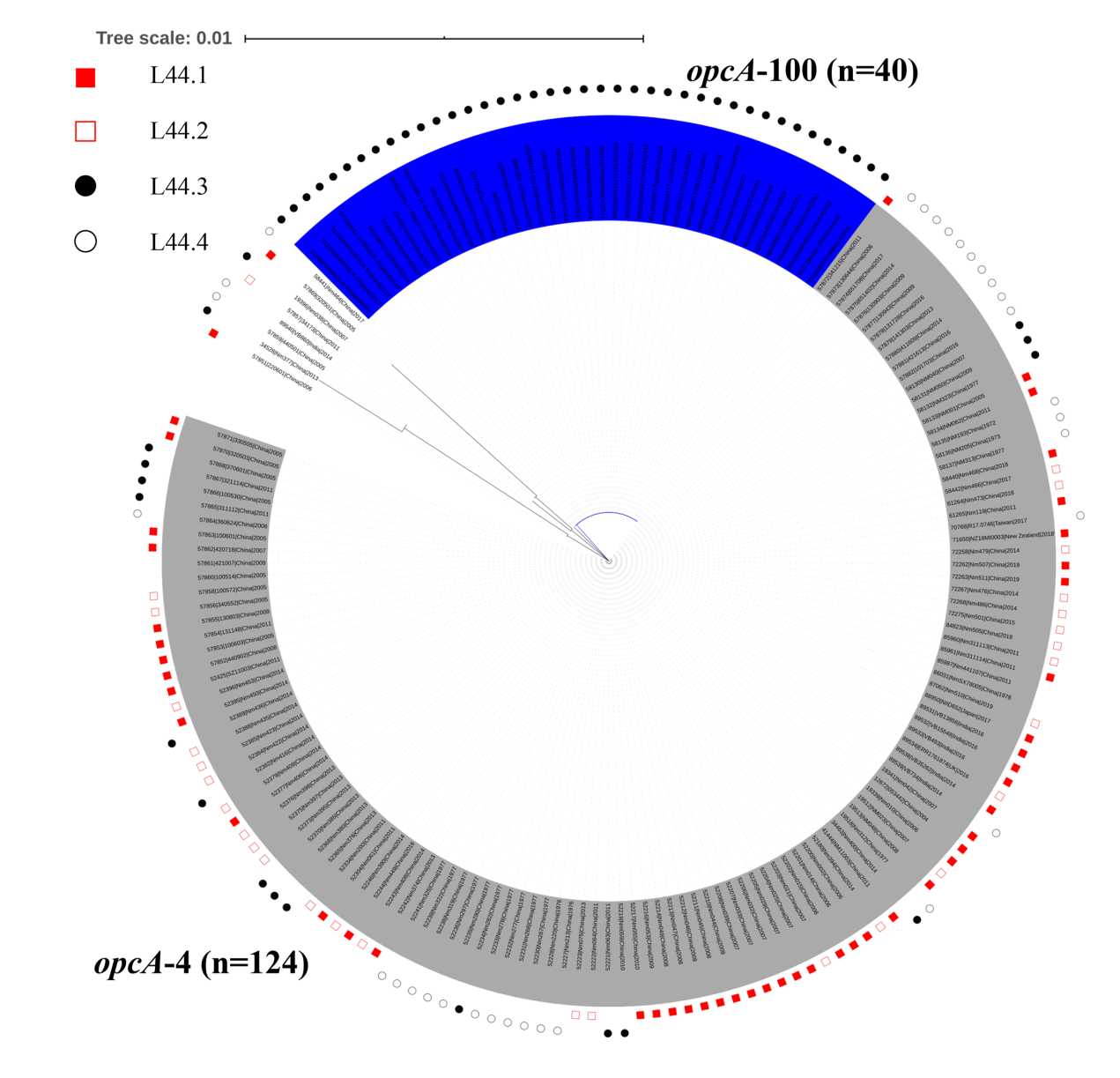

**Figure S7. Phylogenetic analysis of antigen gene sequence variation within the *N. meningitidis* clonal complex 4821 lineage, based on nucleotide sequences of the *opcA* gene.**

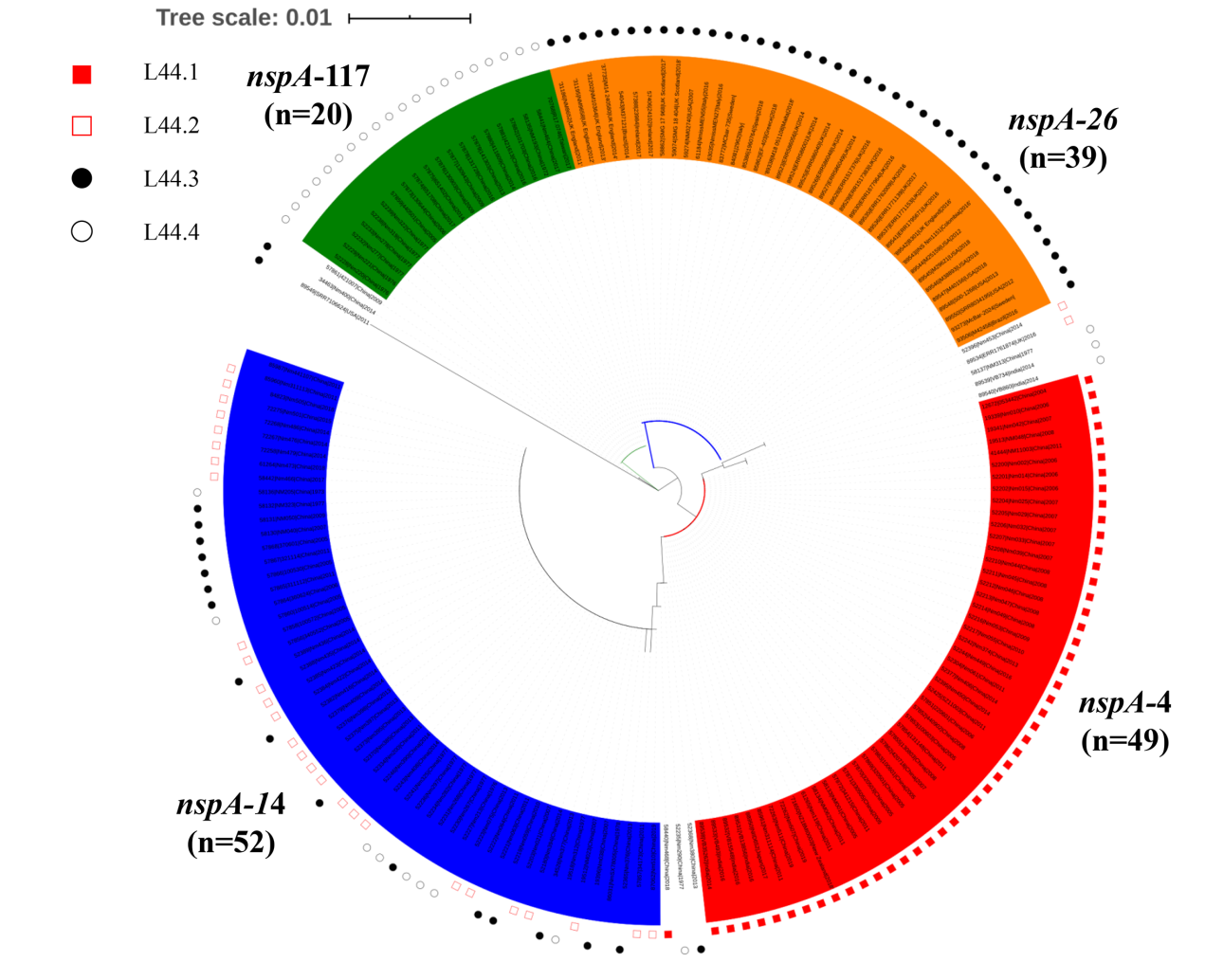

**Figure S8. Phylogenetic analysis of antigen gene sequence variation within the *N. meningitidis* clonal complex 4821 lineage, based on nucleotide sequences of the *nspA* gene.**

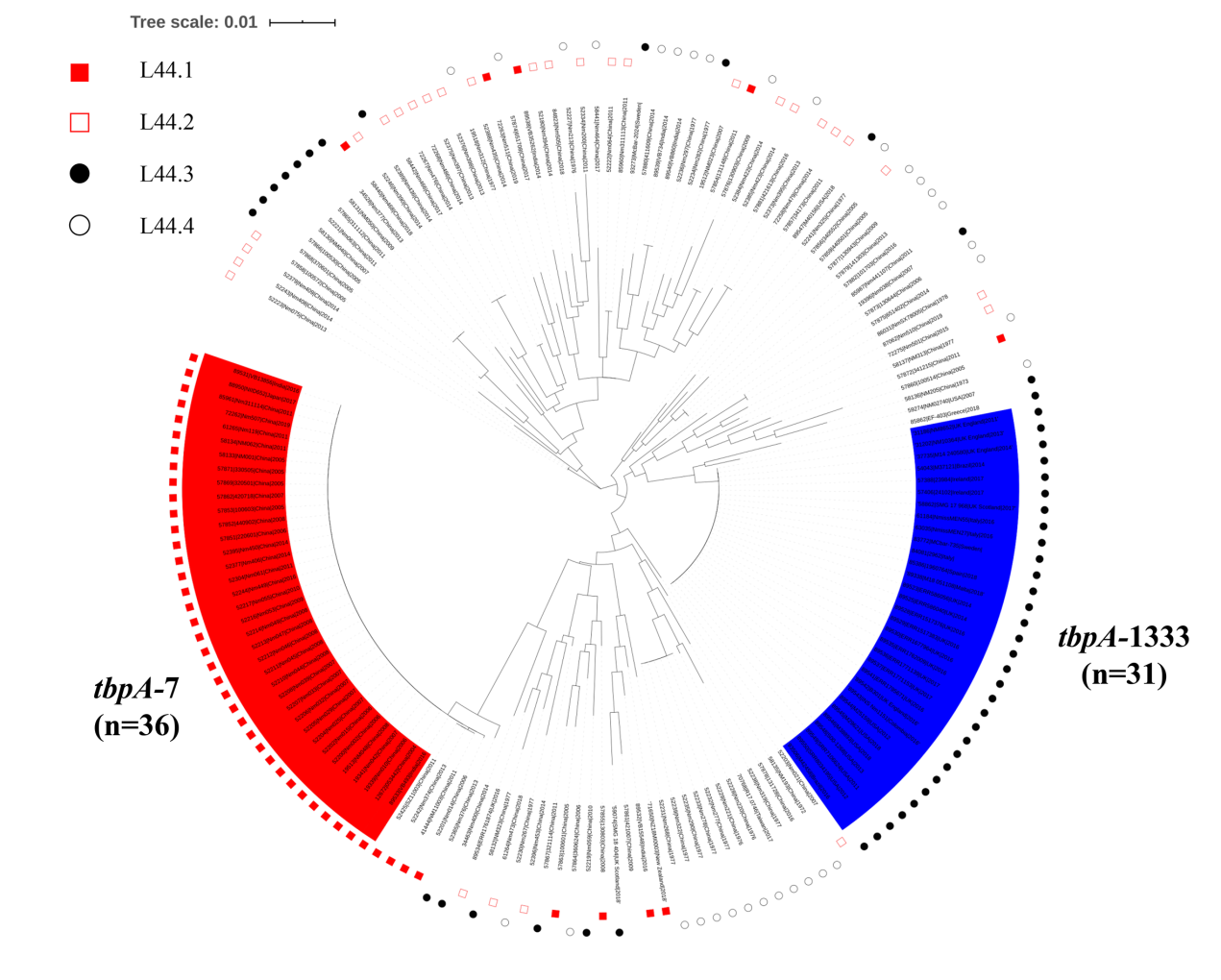

**Figure S9. Phylogenetic analysis of antigen gene sequence variation within the *N. meningitidis* clonal complex 4821 lineage, based on nucleotide sequences of the *tbpA* gene.**

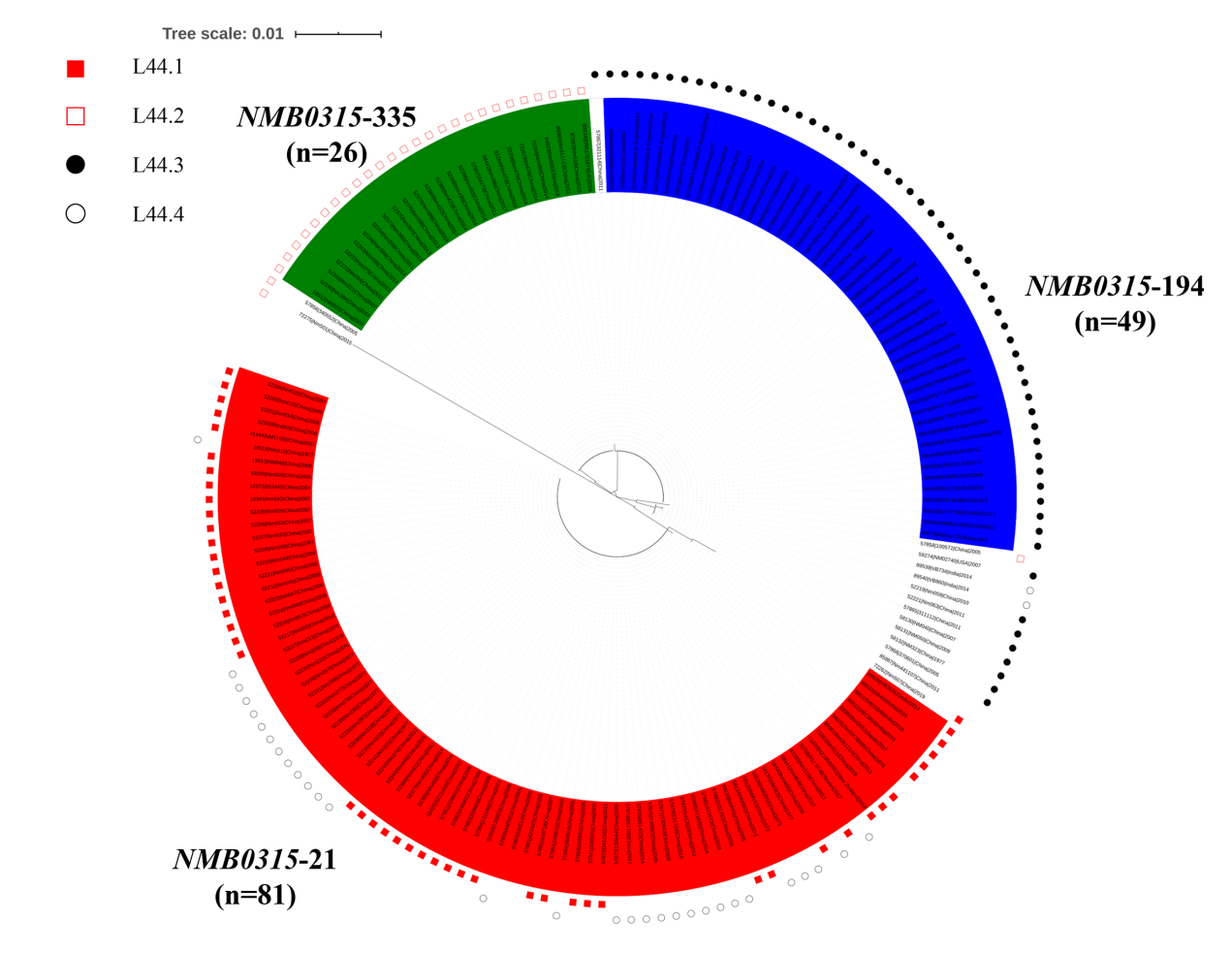

**Figure S10. Phylogenetic analysis of antigen gene sequence variation within the *N. meningitidis* clonal complex 4821 lineage, based on nucleotide sequences of the *NMB0315* gene.**

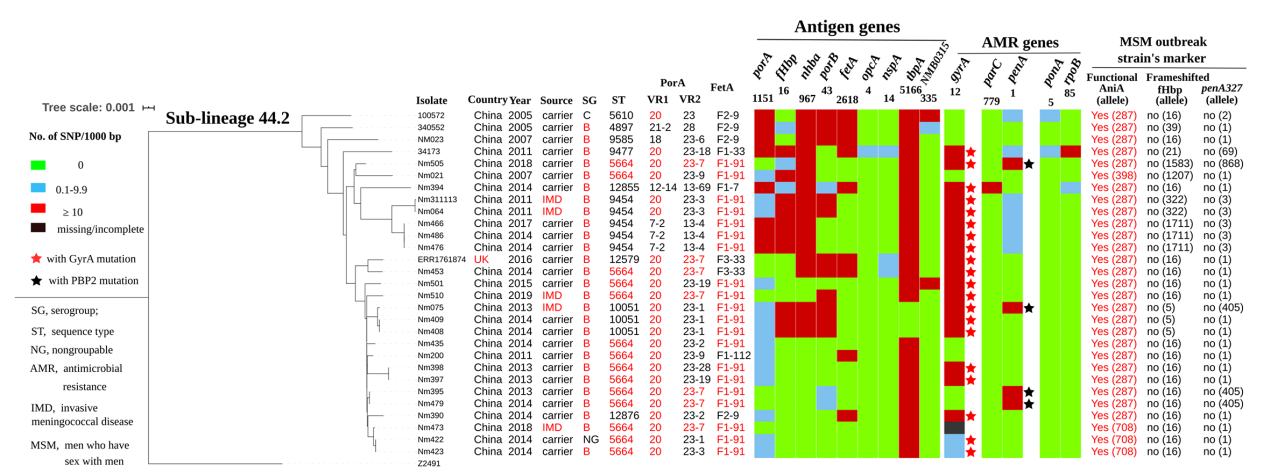

**Figure S11. Genetic diversity of sub-lineage 44.2 isolates.**

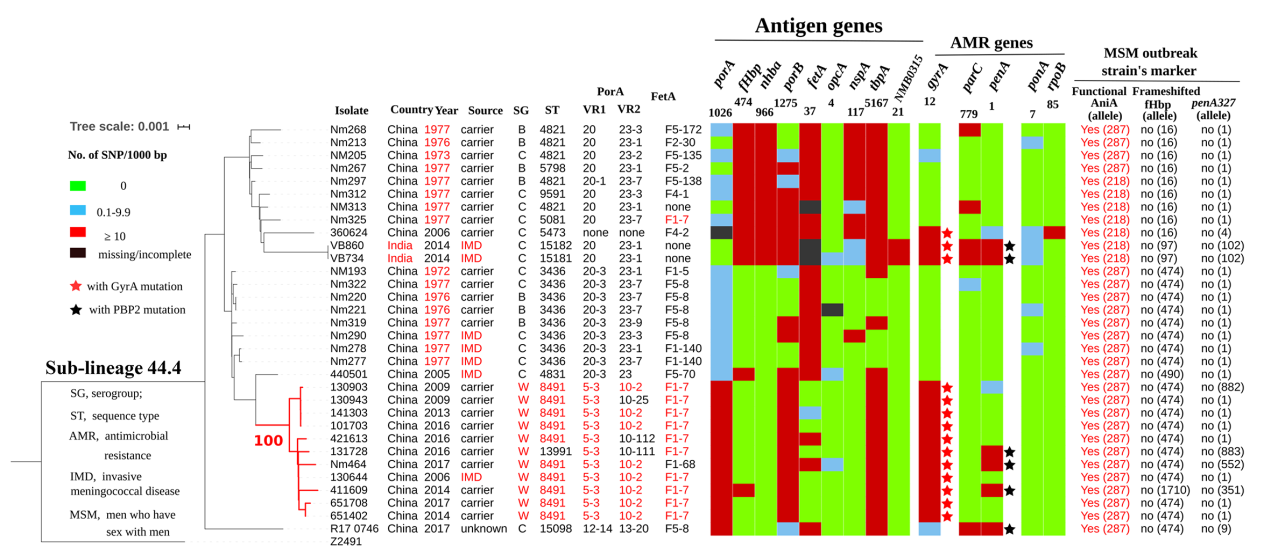

**Figure S12. Genetic diversity of sub-lineage 44.4 isolates.**

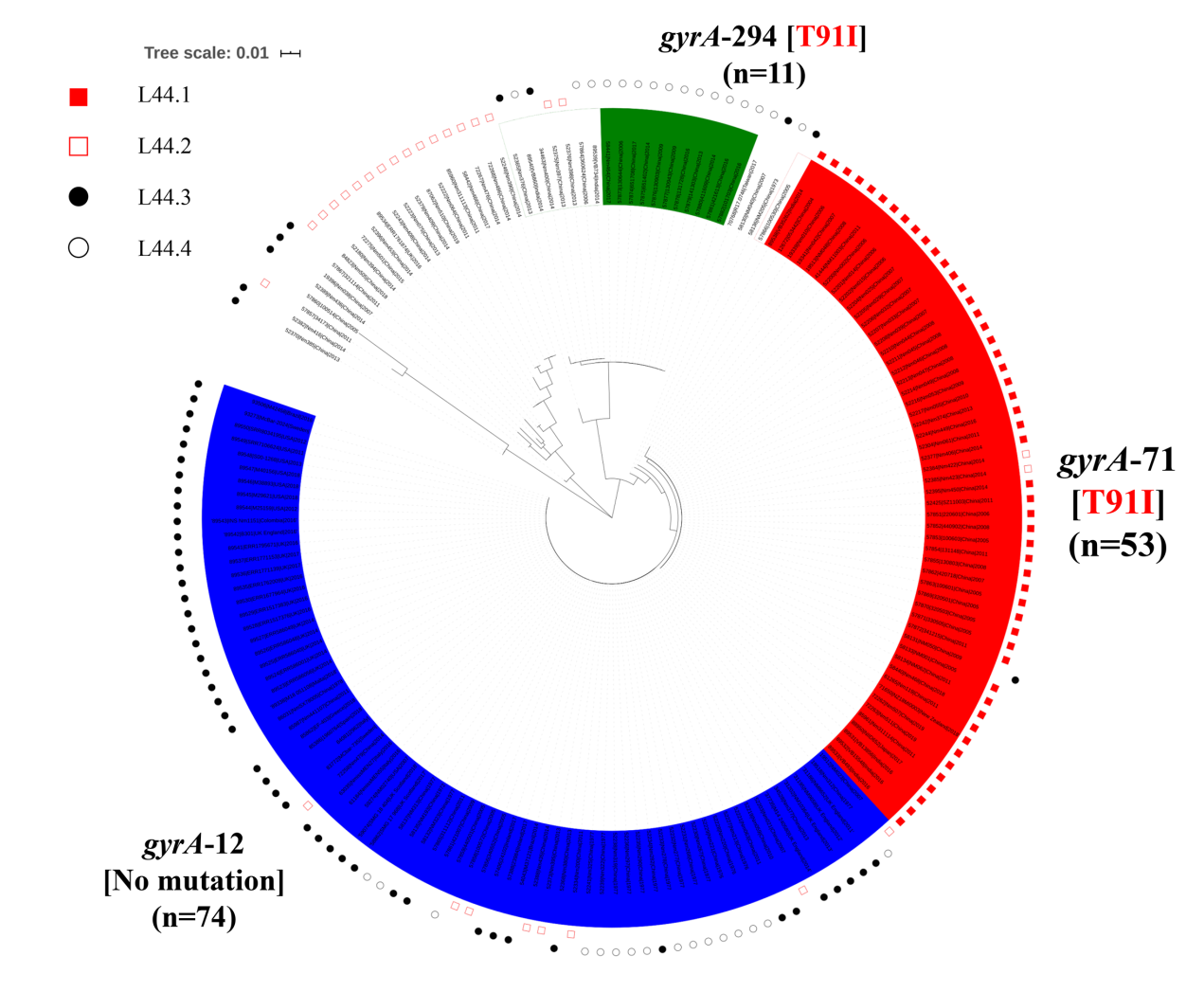

**Figure S13. Phylogenetic analysis of antibiotic resistance gene sequence variation within the *N. meningitidis* clonal complex 4821 lineage, based on nucleotide sequences of the *gyrA* gene.**

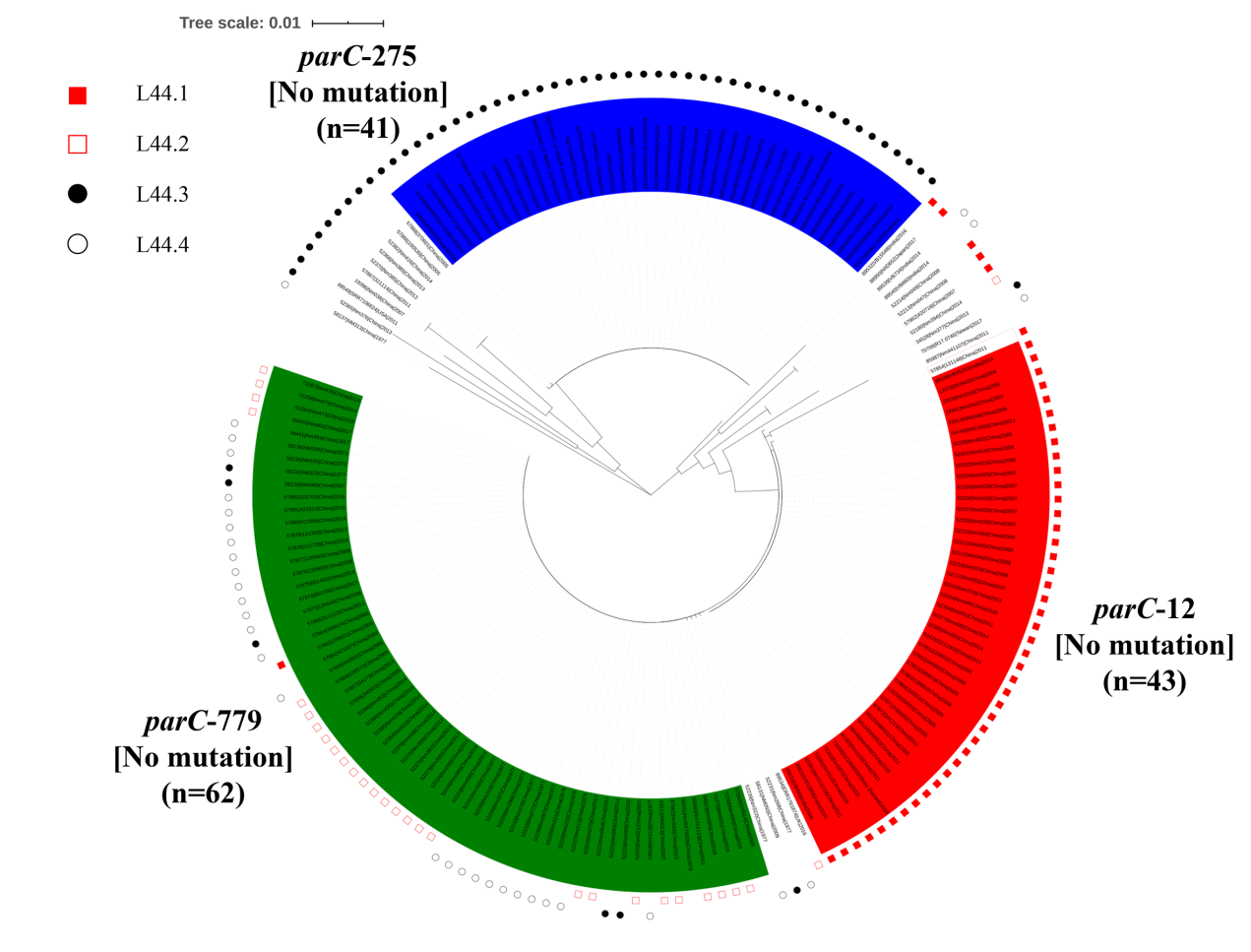

**Figure S14. Phylogenetic analysis of antibiotic resistance gene sequence variation within the *N. meningitidis* clonal complex 4821 lineage, based on nucleotide sequences of the *parC* gene.**

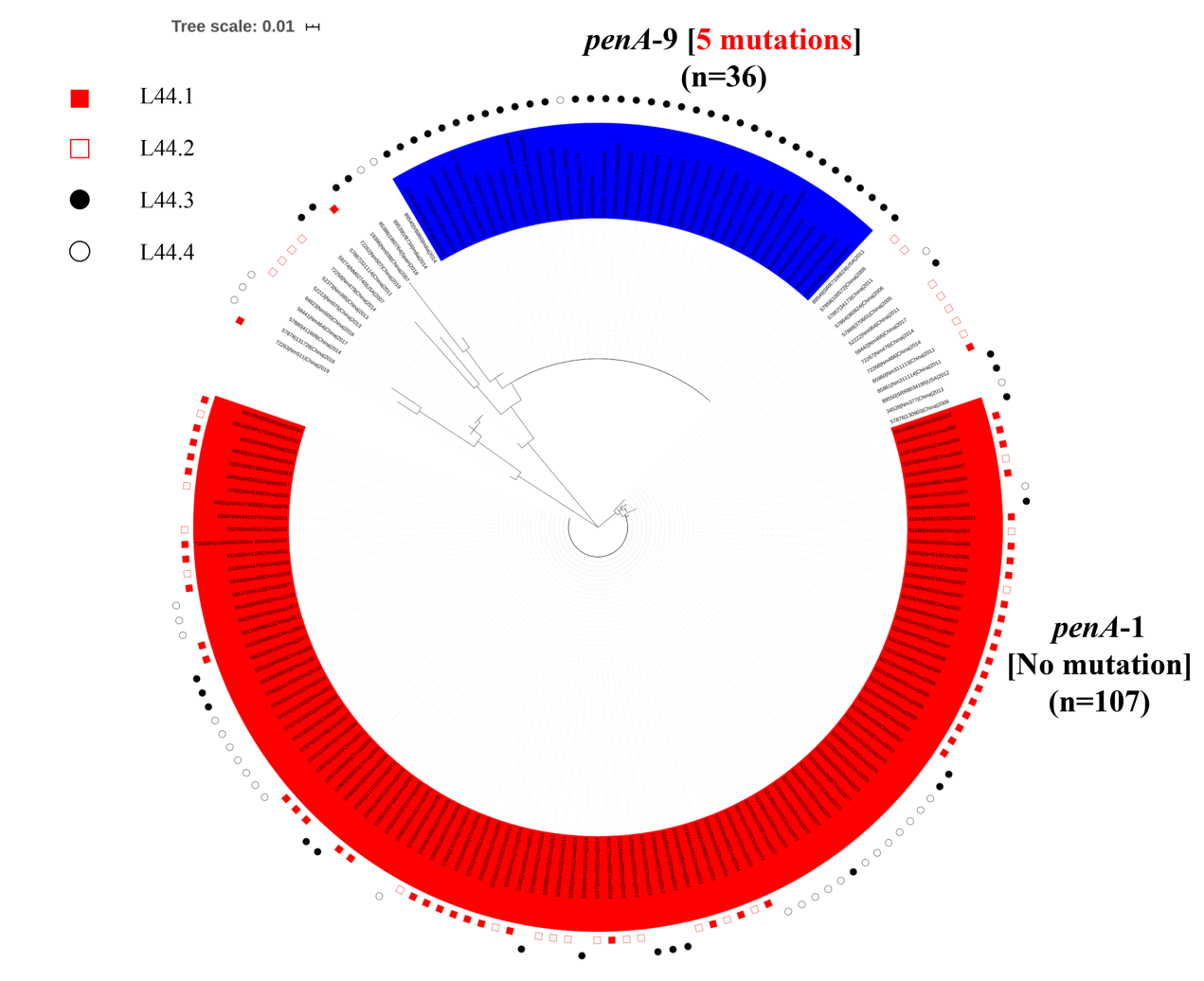

**Figure S15. Phylogenetic analysis of antibiotic resistance gene sequence variation within the *N. meningitidis* clonal complex 4821 lineage, based on nucleotide sequences of the *penA* gene.**

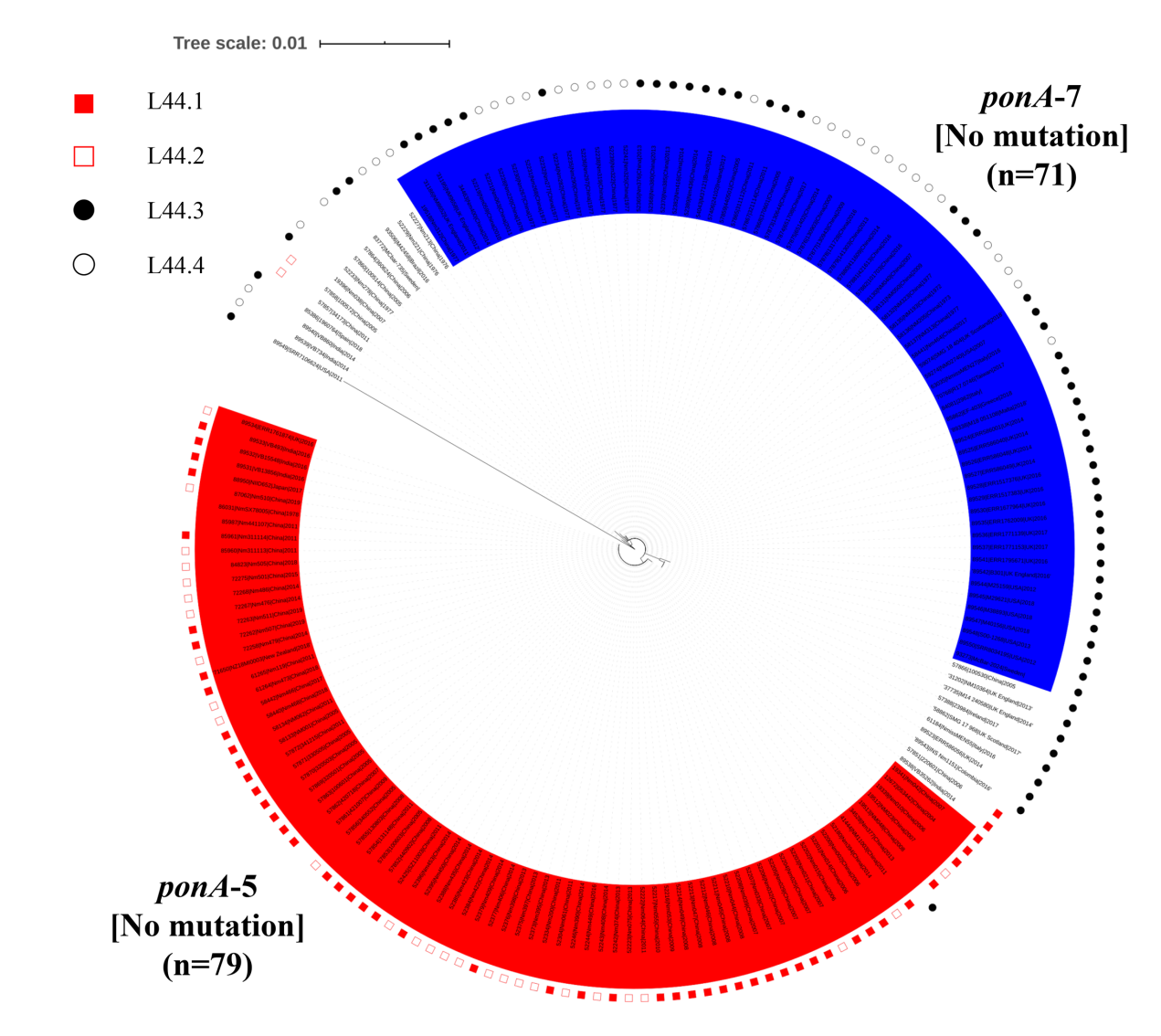

**Figure S16. Phylogenetic analysis of antibiotic resistance gene sequence variation within the *N. meningitidis* clonal complex 4821 lineage, based on nucleotide sequences of the *ponA* gene.**

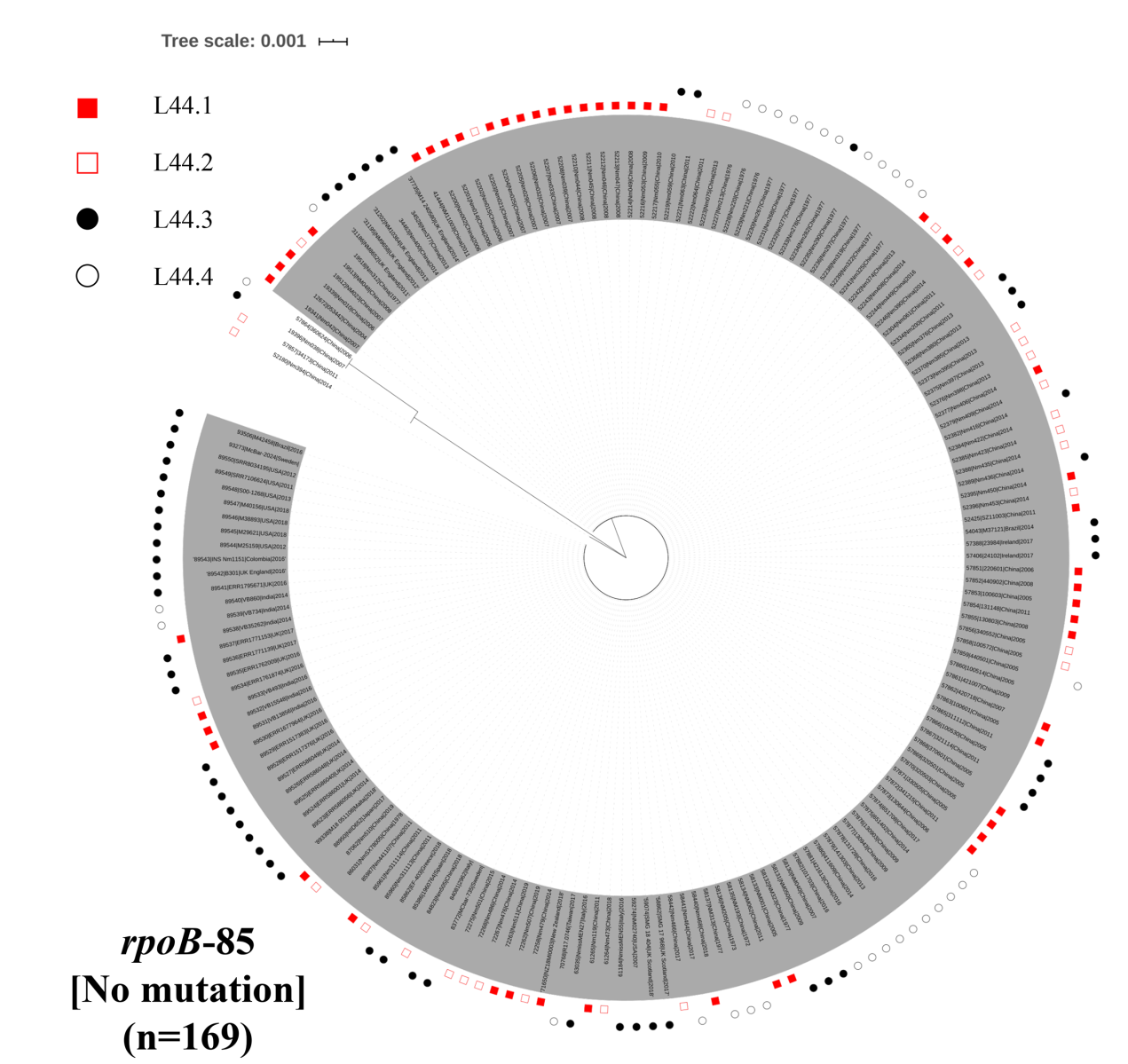

**Figure S17. Phylogenetic analysis of antibiotic resistance gene sequence variation within the *N. meningitidis* clonal complex 4821 lineage, based on nucleotide sequences of the *rpoB* gene.**

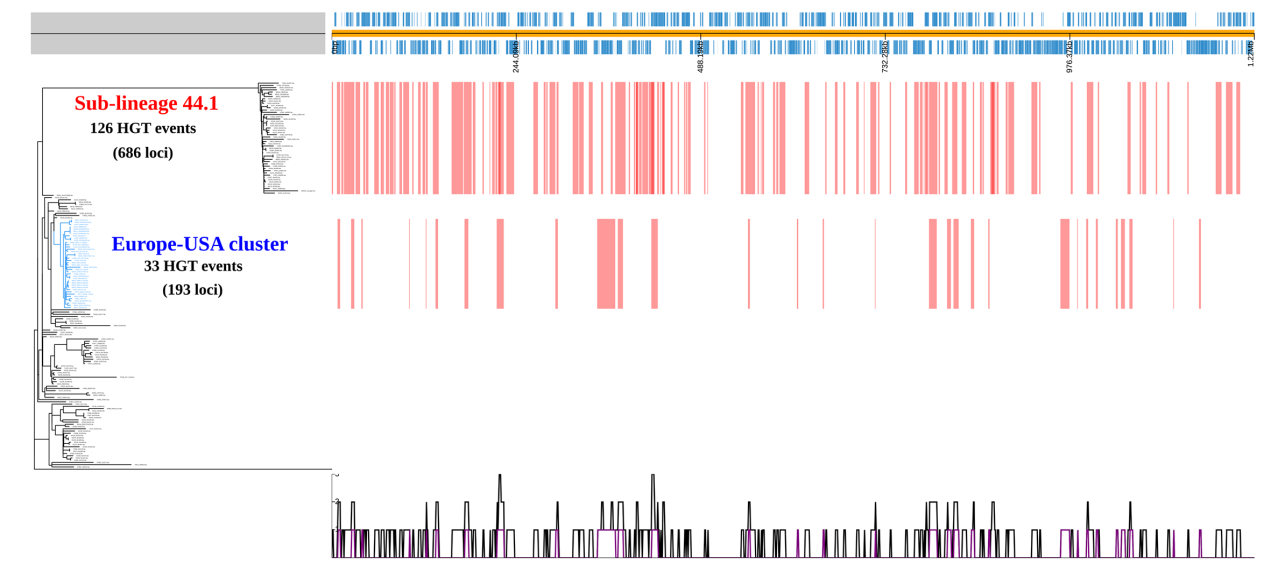

**Figure S18. Horizontal gene transfer (HGT) events predicted by Gubbins.**

Each red vertical line represents a HGT event. A total of 126 HGT events involved with 686 loci in L44.1; 33 events in Europe-USA cluster involved with 193 loci
